# Learning shapes cortical dynamics to enhance integration of relevant sensory input

**DOI:** 10.1101/2021.08.02.454726

**Authors:** Angus Chadwick, Adil Khan, Jasper Poort, Antonin Blot, Sonja Hofer, Thomas Mrsic-Flogel, Maneesh Sahani

**Affiliations:** Gatsby Computational Neuroscience Unit, University College London, London, UK; Sainsbury Wellcome Centre for Neural Circuits and Behaviour, University College London, London, UK; Institute for Adaptive and Neural Computation, University of Edinburgh, UK; Centre for Developmental Neurobiology, King’s College London, London, UK; Department of Physiology, Development and Neuroscience, University of Cambridge, Cambridge, UK; Department of Psychology, University of Cambridge, Cambridge, UK

## Abstract

Adaptive sensory behavior is thought to depend on processing in recurrent cortical circuits, but how dynamics in these circuits shapes the integration and transmission of sensory information is not well understood. Here, we study neural coding in recurrently connected networks of neurons driven by sensory input. We show analytically how information available in the network output varies with the alignment between feedforward input and the integrating modes of the circuit dynamics. In light of this theory, we analyzed neural population activity in the visual cortex of mice that learned to discriminate visual features. We found that over learning, slow patterns of network dynamics realigned to better integrate input relevant to the discrimination task. This realignment of network dynamics could be explained by changes in excitatory-inhibitory connectivity amongst neurons tuned to relevant features. These results suggest that learning tunes the temporal dynamics of cortical circuits to optimally integrate relevant sensory input.

**Highlights:**

- A new theoretical principle links recurrent circuit dynamics to optimal sensory coding
- Predicts that high-SNR input dimensions activate slowly decaying modes of dynamics
- Population dynamics in primary visual cortex realign during learning as predicted
- Stimulus-specific changes in E-I connectivity in recurrent circuits explain realignment

## Introduction

Cortical circuits process sensory information through both feedforward and recurrent synaptic connections (Lamme and Roelfsema, 2000). Feedforward connectivity can filter (Hubel and Wiesel, 1962; LeCun et al., 2015) and propagate (Abeles, 1992; Van Rossum et al., 2002) relevant information, allowing rapid categorization and discrimination of stimuli (Thorpe et al., 1996; Resulaj et al., 2018). However, the majority of synaptic input received by neurons in sensory cortex arises from neighboring cortical cells (Peters et al., 1994; Douglas et al., 1995), and recurrent cortical dynamics exerts a powerful influence on network activity during sensory stimulation (Fiser et al., 2004; Reinhold et al., 2015). The functional role of such recurrent synapses in the integration and transmission of sensory information remains poorly understood.

Many of the stimulus features represented in the spiking output of neurons in primary sensory cortex are already present in the net feedforward input they receive (Lien and Scanziani, 2013). Previous studies have proposed two possible functions of recurrent cortical synapses. First, recurrent synapses may increase the signal-to-noise ratio (SNR) of the relevant sensory features through selective amplification (Douglas et al., 1995; Ben-Yishai et al., 1995; Somers et al., 1995; Murphy and Miller, 2009; Liu et al., 2011; Li et al., 2013; Lien and Scanziani, 2013; Cossell et al., 2015). Second, recurrent synapses may enhance the efficiency of the encoding by suppressing redundant responses in similarly tuned cells (Olshausen and Field, 1996; Lochmann and Deneve, 2011; Chettih and Harvey, 2019). However, although recurrent amplification and competitive suppression can increase the SNR of single-neuron responses and improve coding efficiency respectively, such mechanisms cannot increase the amount of sensory information transmitted through the network beyond the information that the network receives in its input (Cover and Thomas 2006; Seriès et al., 2004; Beck et al., 2011; Kanitscheider et al., 2015; Zylberberg et al., 2017; Huang et al., 2020).

Recent studies have shown that visual features such as orientation become easier to decode from both single-cell and population responses in primary visual cortex (V1) when mice and monkeys learn to associate them with behavioral contingencies (Poort et al., 2015; Khan et al., 2018; Jurjut et al., 2017; Yan et al., 2014). This apparent improvement in representation is accompanied by changes in functional interactions amongst excitatory and inhibitory cell types within the local circuit (Khan et al., 2018). Since changes in recurrent amplification or competitive suppression cannot increase the total available information, it remains unclear how changes in the local circuit could generate the observed improvements.

Here, we ask whether improvements in stimulus decodability over learning could arise through selective temporal integration of relevant feedforward sensory input. We first show analytically how the output of a network can be tuned to optimally discriminate pairs of input stimuli by matching its recurrent dynamics to their sensory input statistics. In particular, we show that a stimulus decoder applied to network output performs best if the dimension of network input with greatest SNR activates a pattern of recurrent network dynamics that decays slowly. We then study how the dynamical properties of neural circuits in mouse V1 change as animals learn to discriminate visual stimuli. Using a dynamical systems model fit to experimental data (Khan et al., 2018), we find that slowly decaying patterns in the recurrent dynamics became better aligned with high-SNR sensory input over learning. Finally, we analyze circuit models with excitatory and inhibitory neurons to explore how this alignment might arise through changes in the circuit. We find that stimulus-specific changes in connectivity between excitatory and inhibitory neurons increase the alignment of recurrent dynamics with sensory input as observed experimentally. These connectivity changes predict changes in stimulus tuning within the model, which we find to be recapitulated in the experimental data. Our findings suggest a critical role for cortical dynamics in selective temporal integration of relevant sensory information.

## Results

### Sensory discrimination relies on temporal integration of optimally weighted sensory input

We first asked how the dynamical properties of a recurrent network influence its capacity to discriminate sensory inputs. The scenario we considered had one of two possible stimuli appear for the duration of a trial. Each stimulus generated an input to each neuron in the network with constant mean corrupted by additive, temporally uncorrelated, Gaussian noise (this approximates the net feedforward synaptic input a neuron receives from a large number of upstream neurons, see Stein, 1967; Capocelli and Ricciardi, 1971; Lansky, 1984). To determine how these inputs should be integrated for optimal discrimination performance, we adopted a signal processing perspective (see Supplementary Mathematical Note).

Two noisy stimuli can be optimally discriminated from the instantaneous sensory input to the network by taking a one-dimensional linear combination of the inputs to different neurons (Figure 1A, B) weighted according to the “linear discriminant". This is the linear combination of inputs that achieves the best compromise between separating the mean inputs under the two stimuli and avoiding projected noise (Figure 1D, black dashed arrow). Writing **u**(*t*) for a vector collecting the inputs to all neurons at time t, the linear discriminant is a vector w of the same dimension such that the projected input vector *d*(*t*) = **w · u**(*t*) has the greatest possible signal-to-noise ratio SNR_input_(**w**) for the discrimination of the two stimuli (Figure 1B, D). Then, to discriminate stimuli over a window of duration T, the optimal strategy is simply to integrate the linear discriminant projection across the time window (Figure 1C), yielding an output with 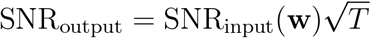 (Figure 1E, F).

**Figure 1.**
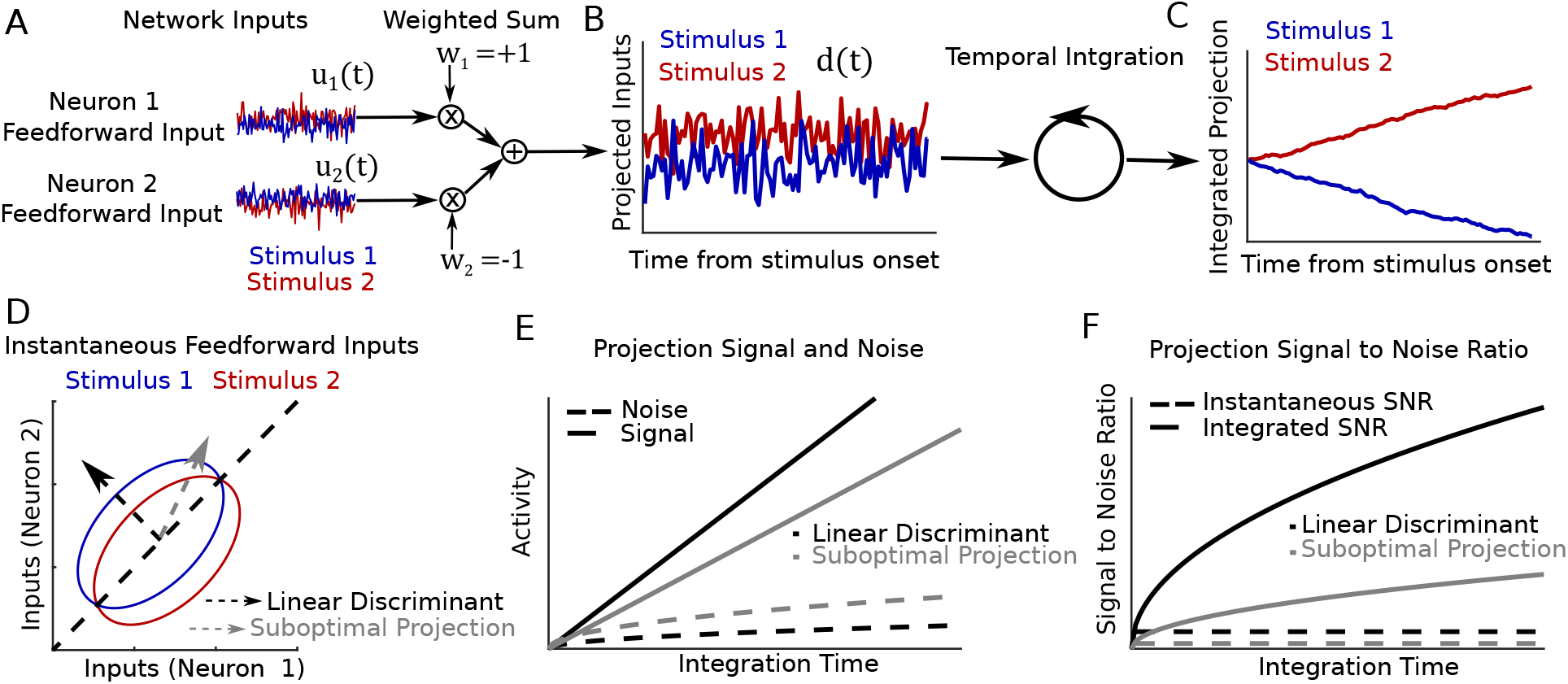
Stimulus discrimination performance depends on temporal integration of weighted sensory input. A: Feedforward inputs to a two-neuron network, shown for two different stimuli (red and blue). B: A weighted sum (linear projection) of the instantaneous inputs shown in A. C: The temporally integrated input projection for each stimulus (cumulative sum of projected inputs shown in B). D: Distributions of instantaneous feedforward input for each of the two stimuli (colored ellipses), their optimal linear discriminant (dashed black arrow), and a second suboptimal projection (dashed gray arrow). E: The signal (difference in mean; solid lines) and noise (standard deviation; dashed lines) of activity following linear projection and temporal integration, shown for the two projections in D. F: The instantaneous (dashed) and temporally integrated (solid) signal to noise ratio of these two projections.

These results demonstrate that a network can best generate distinct activity patterns in response to two different continuous stimuli if it temporally integrates the input stimuli weighted according to their projection onto an optimal linear discriminant.

### Recurrent networks enhance sensory discrimination by alignment of slowly decaying dynamical modes with optimal sensory input

How might this optimal discrimination function be achieved using a recurrent network? To address this, we considered how noisy stimulus input is filtered through the recurrent network dynamics. A core feature of recurrent networks is their capacity to generate multiple distinct activity patterns, which may unfold with different dynamical time constants within the network’s high-dimensional activity space (Rabinovich et al., 2006; Miller, 2016; Sussillo et al., 2014). We asked if these different time constants of network dynamics could allow a network to act as an optimal integrator of sensory input by providing windows of temporal integration over the optimal input discriminant (Goldman et al., 2009a).

For networks that settle into a steady pattern of firing rates when driven by a constant input (Figure 2A, C), the behavior of small fluctuations around that input-driven fixed point can be approximated with a linear dynamical system (Figure 2B). The dynamics of this linearized network are described by a set of dynamical “modes", each of which associates a time constant *τ* with a unique pattern of network activation **m** (Figure 2B). The activation pattern **m** is a vector describing a particular deviation of network activity from the fixed point, with elements equal to the relative deviation of each neuron, while *τ* determines the time taken for an activity fluctuation along **m** to decay back towards the fixed point through the network dynamics. In particular, when network activity is perturbed away from its input-driven fixed point along any direction, the ensuing population activity trajectory projected onto any given mode’s **m** decays as an exponential function with the corresponding time constant *τ* (Figure 2B, C). Moreover, when the network is driven by a stimulus input with continuously fluctuating noise as considered here (Figure 1A), population activity projected onto any mode’s **m** behaves as a leaky integrator, with each mode independently aggregating inputs that fall along its activation pattern with an integration window of duration *τ* (Figure 2D, E). In the discrimination task, input associated with one of the two possible stimuli drives the network on any given trial (Figure 1A, D, Figure 2D). In this case, provided that the two stimulus-driven fixed points are sufficiently close to fall within the domain of network linearization (Figure 2E, F), the SNR of network output projected onto any single mode’s **m** following network integration matches the signal processing solution above, with 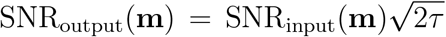 (Figure 2I, J). Thus, a recurrent network can achieve the optimal strategy for stimulus decoding (Figure 1) if its recurrent connectivity gives rise to a dynamical mode with activation pattern **m** that is aligned to the input linear discriminant **w** (i.e., **m** = **w**) and decay time constant *τ* that is longer than the stimulus window *T* (as in Figure 2E, F; panels G, H show suboptimal integration). In other words, the recurrent dynamics are optimized for discrimination of a pair of input stimuli with linear discriminant **w** if fluctuations of network activity along **w** decay slowly.

**Figure 2.**
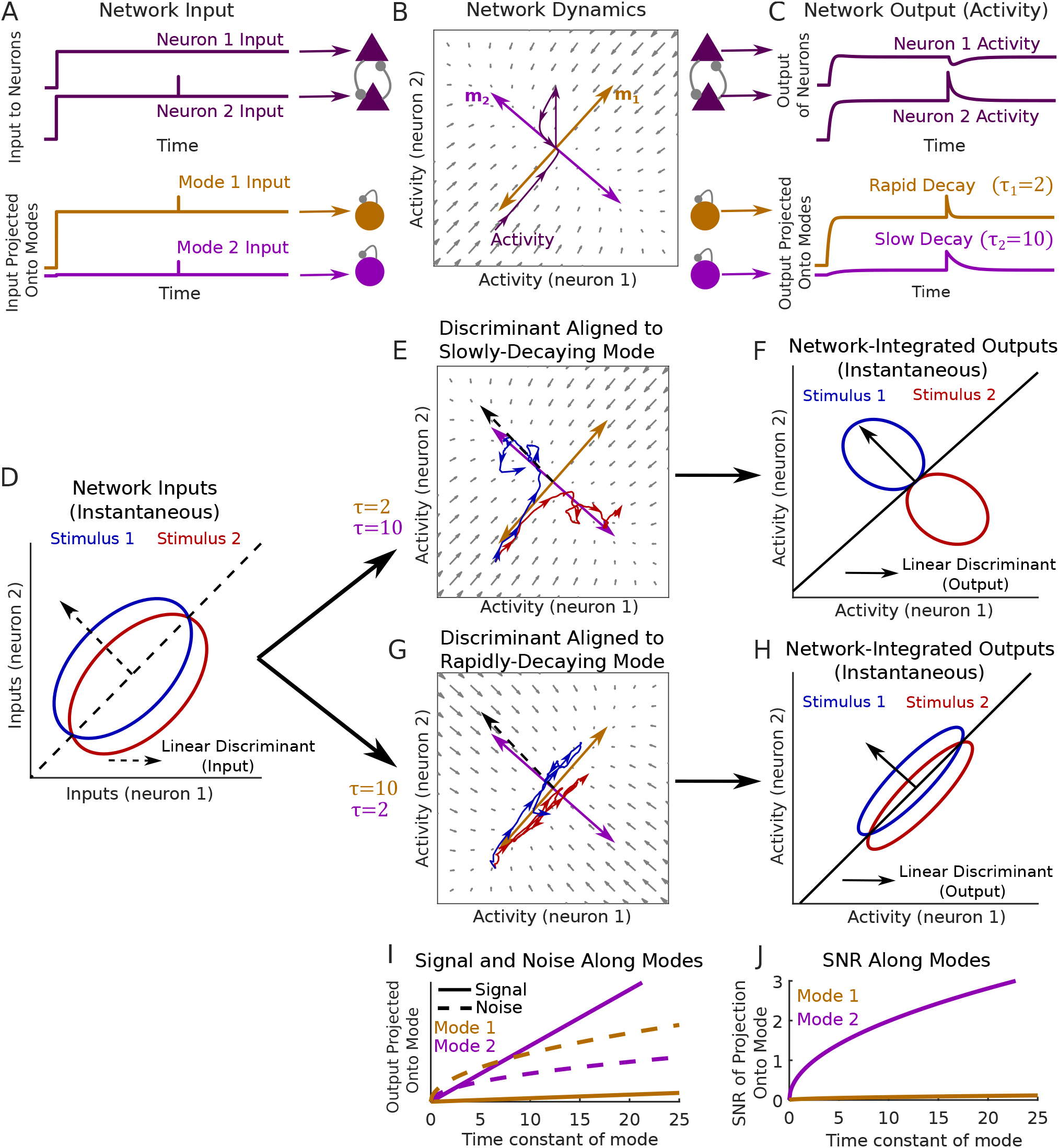
Alignment of dynamical modes with feedforward input determines sensory discrimination performance. A-C: Illustration of a two-neuron network receiving feedforward input and generating an output activity pattern with rapidly and slowly decaying dynamical modes (brown and light purple). A: (Top). Constant input to each neuron, and a small input perturbation to neuron 2. (Bottom) The same input shown following projection onto the two modes of network dynamics. B: Illustration of network dynamics. Gray arrows depict the dynamical flow of network activity from a given state when input is held at the constant level shown in A. Light purple and brown arrows depict modes’ activation patterns m. The trajectory of neural activity in response to the input in A is shown in dark purple. The input perturbation to neuron 2 generates a dynamical response along both modes, each decaying with a different time constant *τ* C: Network output shown for each neuron and along each mode. Single-neuron responses exhibit complex and heterogeneous timecourses, but the network response projected onto any mode exhibits a simple exponential decay. D: Distributions of instantaneous feedforward input under two different stimuli (red and blue ellipses), as in Figure 1A, D (note that inputs have time-varying noise). E: A network with a slowly decaying mode aligned to the input linear discriminant. Blue and red traces show example trajectories of network output when the network is driven by a single-trial input from each of the two stimulus distributions. F: Distributions of instantaneous network output at equilibrium under each stimulus. G, H: As in E, F but with a rapidly decaying mode aligned to the input linear discriminant. I: Signal and noise of instantaneous network output along each mode, as a function of the mode’s time constant. J: Signal-to-noise ratio of instantaneous network output along each mode.

Biological neural networks may exhibit complex “non-normal” dynamics, including rapid “balanced amplification” and temporally extended “functionally-feedforward” activation (Ganguli et al., 2008; Murphy and Miller, 2009; Goldman, 2009b). In functionally-feedforward networks, activation of one group of neurons causes subsequent activation of other neuron groups, leading to transient activity sequences whose lifetime exceeds the decay time of any individual mode (Goldman, 2009b). We asked whether these non-normal dynamics might yield further mechanisms for optimizing stimulus discrimination. We found analytically that the discrimination performance of a network depends on the geometry of its modes’ activation patterns (Supplementary Figure 1A, B). When these are orthogonal, corresponding to “normal” networks, response information is maximized when the most slowly decaying mode has activation pattern aligned to the input linear discriminant (Figure 2E, Supplementary Figure 1A, B). Analyzing “non-normal” networks, we found that response information further improves when multiple modes have their activation patterns aligned with the input linear discriminant (Supplementary Figure 1A, B). These improvements arise through functionally-feedforward dynamics, which increase the total window of network integration relative to the decay time constants of the individual modes (Supplementary Figure 1A, C-H) (Ganguli et al., 2008; Goldman, 2009b).

Taken together, our findings demonstrate that recurrent networks maximize their capacity to discriminate sensory inputs when they align one or more slowly decaying modes of dynamics with the optimal input discriminant. We reasoned that such a mechanism may underlie improvements in cortical representations for relevant stimuli over learning (Poort et al., 2015; Khan et al., 2018).

### Learning reorganizes cortical networks to enhance integration of relevant sensory input

With this description of recurrent processing in mind, we examined the effects of learning on cortical dynamics and sensory representations. We analyzed the activity of neuronal populations in primary visual cortex of head-fixed mice as they learned to perform a visual discrimination task within a virtual reality environment. Over a period of 7-9 days, mice learned to selectively lick a reward spout in a virtual corridor lined with vertical but not angled stripes (Figure 3A, B). The responses of the same populations of neurons to these stimuli were measured before and after learning using chronic two-photon calcium imaging. Learning led to an improvement in the linear discriminability of these two stimuli based on instantaneous population responses (Figure 3E right, p = 0.035, one-sided sign test on pre- vs post-learning discriminability, see Methods for details). Given that instantaneous sharpening or amplification of sensory input by the V1 circuit cannot increase response information (Cover and Thomas 2006; Seriès et al., 2004; Beck et al., 2011), we hypothesized that such improvements could arise via either 1) an increase in sensory information provided through external input to the circuit (i.e., an increase in SNR_input_(**w**) caused by changes in upstream processing) or 2) a reorganization of local circuit dynamics to enhance temporal integration of sensory input (Figures 1, 2).

**Figure 3.**
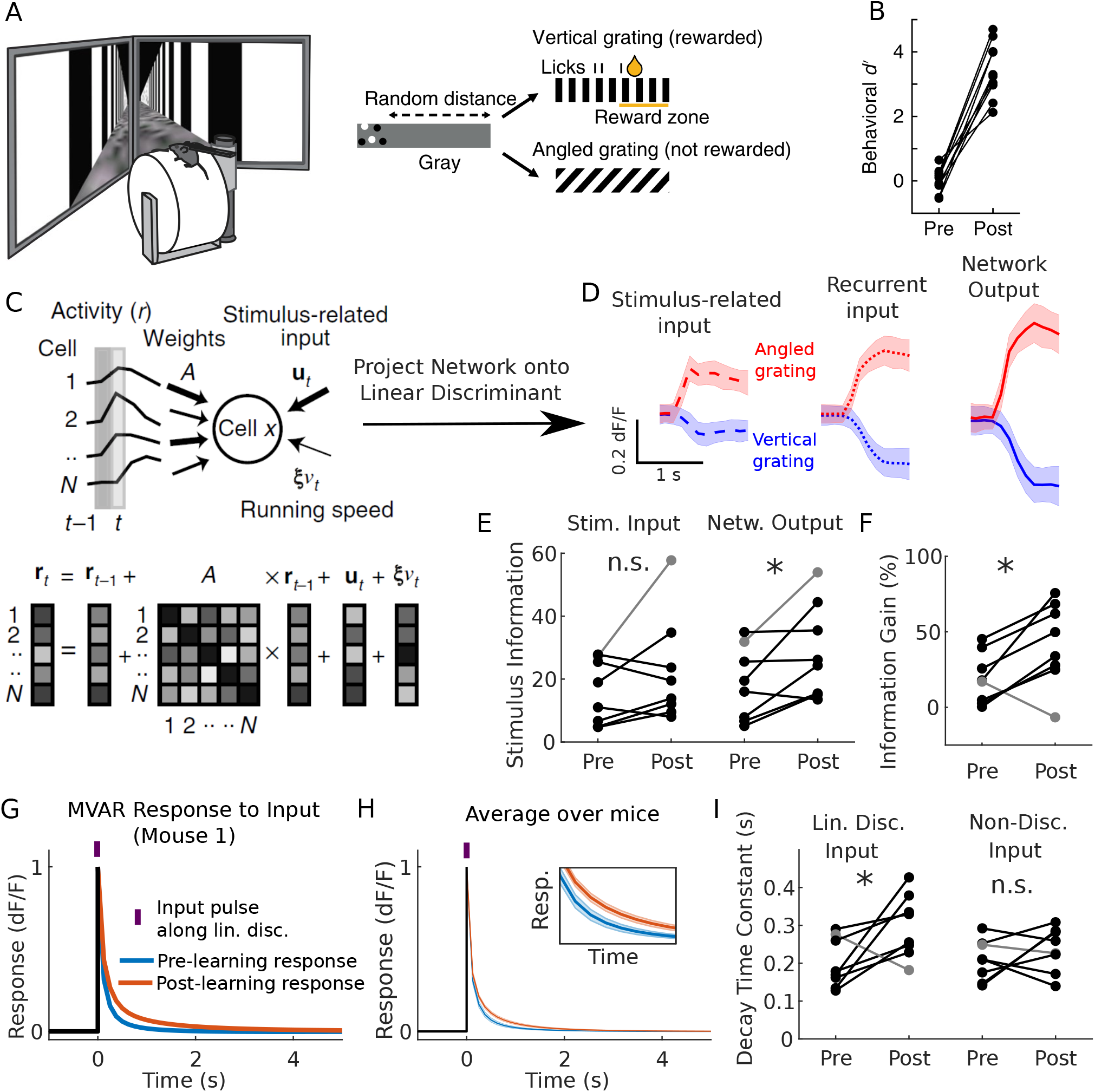
Changes in V1 population dynamics over learning selectively enhance temporal integration of relevant sensory input. A: Visual discrimination task. B: Behavioral performance of each mouse pre- vs post-learning. C: Schematic describing MVAR model fit to imaged population activity. The MVAR model fits variability in single-trial responses of each cell by estimating the contribution of stimulus-locked input, recurrent input from the local cell population, and running speed. D: The inferred stimulus-related and recurrent input and the imaged network output, each projected onto the optimal linear discriminant (mean ± standard deviation over trials for one mouse post-learning). E: Information in MVAR stimulus-related input and network output for each mouse pre- vs post-learning (gray line delineates a particular mouse whose improvements occurred through enhanced stimulus-related input). F: MVAR input-output information gain, pre- vs post-learning for each mouse. G: Simulated response of the MVAR model to a synthetic pulse of input aligned to the linear discriminant, pre- and post-learning for one mouse. H: As in G, showing mean±sem over mice. Inset shows zoomed in traces. I: Left: The decay time constant of responses in G and H for each mouse, pre- vs post-learning. Right: The decay time constants for a second input pattern that carries no information about stimulus identity.

Distinguishing these hypotheses requires a complete characterization of the dynamics of the imaged circuit and the sensory input it receives before and after learning. As it is not currently possible to achieve this experimentally, we sought to infer the recurrent dynamics and stimulus inputs which best accounted for the coordinated activity patterns of the imaged circuit using a statistical model fit to the data. To this end, we examined a multivariate autoregressive (MVAR) linear dynamical system model we had previously fit to population activity imaged before or after learning (Khan et al., 2018). The MVAR model predicts the activity of each cell at imaging frame t based on 1) recurrent input from all imaged cells at time step t-1, with stimulus-independent weights; 2) a time-varying stimulus-dependent input, locked to stimulus onset and the same for all trials with a given stimulus; and 3) the running speed of the animal at time t (Figure 3C). Imaged responses in the population covaried in time and across trials, in a way that could not be explained by changes in the stimulus or changes in running behavior (Khan et al., 2018). The model depended on the recurrent interaction term to capture such “noise” covariance, and so once the model was fit to data these weights were effectively determined by the structure of observed trial-by-trial variability. Conversely, the stimulus-dependent trial-invariant terms were determined during fitting so that the input signals, once fed through the recurrent terms of the model, captured the trial-averaged response profiles. Any remaining trial-by-trial variability in the data was assigned to a residual term (see Methods and Khan et al., 2018 for a more detailed discussion of the MVAR model and its validation on the present dataset). Given this characterization of the imaged responses in terms of stimulus-related input and recurrent interactions (Figure 3D), we then sought to determine the respective contributions of these components to the improvements in response information over learning (Figure 3E right).

To assess whether input information increased over learning, we computed the linear discriminability of stimuli based on the stimulus-related input inferred by the MVAR model, assigning model residuals to noise in this input (Figure 3D, left). Information contained in this input did not increase (p>0.36, one-sided sign test on linear discriminability pre- vs post-learning over all mice; Figure 3E, left). However, there was an increase with learning in the gain of output-to-input information for 7/8 mice (Figure 3E, F, p=0.035, one-sided sign test on relative percentage difference between MVAR input and output information). Thus, the MVAR model ascribed improvements in population response information to learning-related changes in recurrent interactions acting on stimulus-related input that was itself unchanged in information content.

If these recurrent changes acted to improve temporal integration, then the network response to an input pattern aligned with the linear discriminant should be observed to decay more slowly after learning than before. Indeed, the MVAR response to a pulse of such input decayed more slowly after learning for all mice in which improvements in response information were attributed to recurrent dynamics (p=0.035, one-sided sign test on all mice, Figure 3G-I). Moreover, when this analysis was repeated for a second input pattern that was orthogonal to the input discriminant, the decay time did not change over learning (p=0.64, one-sided sign test, Figure 3I, right). Thus, learning induced changes in temporal integration which were selective for task-relevant sensory input.

Enhanced temporal integration could arise through changes in the interaction weights or the stimulus-related input (for example, if stimulus input realigned to drive more slowly decaying network activity patterns). To distinguish between these possibilities, we refit the MVAR model with either interaction weights or stimulus-related input constrained to remain fixed over learning (see Methods). Changes in temporal integration did not occur when interaction weights were fixed (p=0.36, one-sided sign test) but persisted when stimulus-related input was fixed (p=0.004, one-sided sign test, Supplementary Figure 2A, B). This suggested that the improvements relied on changes in interaction weights but not stimulus input.

Taken together, these findings suggest that stimulus information in network responses improved over learning through changes in recurrent dynamics that selectively enhanced temporal integration of task-relevant sensory input.

### Enhanced integration depends on realignment of slowly decaying modes with sensory input

Altered recurrence could selectively enhance temporal integration of relevant sensory input in two ways. First, it could lengthen the decay time constants of those modes whose activation patterns are already best aligned with the input linear discriminant (‘dynamical slowing hypothesis’, Figure 4A, B). Alternatively, it could realign the activation patterns of existing slowly decaying modes towards that discriminant (‘dynamical realignment hypothesis’, Figure 4C).

**Figure 4.**
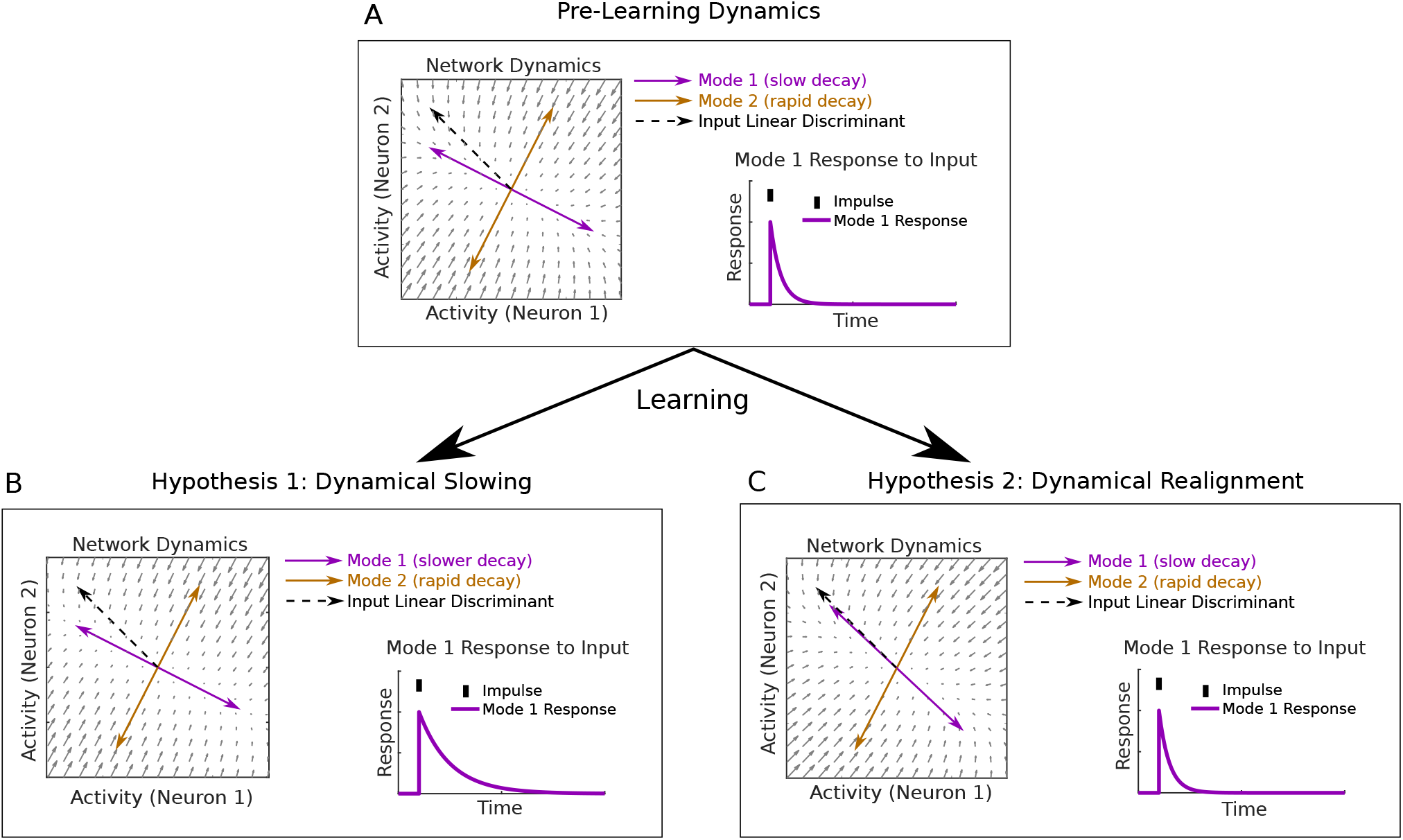
Improvements in temporal integration of relevant sensory input could arise from either slowing or realignment of dynamical modes. A: Example of pre-learning dynamics for a two-neuron network. B: According to the dynamical slowing hypothesis, modes whose activation patterns are best aligned with the input linear discriminant extend their decay time constants over learning, leading to longer timescales of integration over the relevant input patterns. C: In the dynamical realignment hypothesis, modes which decay most slowly become better-aligned to the input linear discriminant over learning.

To distinguish between these two hypotheses, we computed modes of network dynamics and their time constants from the pre- and post-learning MVAR interaction weight matrices. For each mode, we computed the proportion of stimulus-related input information that fell along its activation pattern (its “normalized input SNR”, SNR_norm_(**m**) = SNR_input_(**m**)/SNR_input_(**w**), which is maximized when the mode is aligned to the input linear discriminant). The dynamical slowing hypothesis predicts that the time constants of modes with high input SNR should increase (Figure 4A, B). However, the time constants of modes did not change significantly over learning, either across all modes (p>0.79, one-sided Wilcoxon rank sum test on pre- vs post-learning time constants for all modes pooled across animals) or the subset modes with high input SNR (Figure 5A, B). In contrast, the dynamical realignment hypothesis predicts that the normalized input SNRs of slowly decaying modes should increase (Figure 4A, C). This prediction was borne out by a striking increase over learning in normalized input SNR (p=0.03, one-sided Wilcoxon rank sum test on all modes pooled across animals pre- vs post-learning) which was most pronounced for modes with time constants of ∼700-1000 ms (Figure 5C, D). The increase in normalized input SNR occurred for 7/8 mice (p=0.035, one-sided sign test on average over modes within each mouse pre- vs post-learning, Supplementary Fig 3A), while time constants increased for only 3/8 mice (p=0.86, one-sided sign test on average over modes within each mouse pre- vs post-learning, Supplementary Fig 3B). Examining the joint distribution of the time constants and normalized input SNRs of modes before and after learning (Figure 5E, F), we found a fall in the number of slowly decaying modes with low input SNR matched by an increase in the number with similar decay time constants but high input SNR. These changes are consistent with a realignment of slowly decaying modes towards the input linear discriminant.

**Figure 5.**
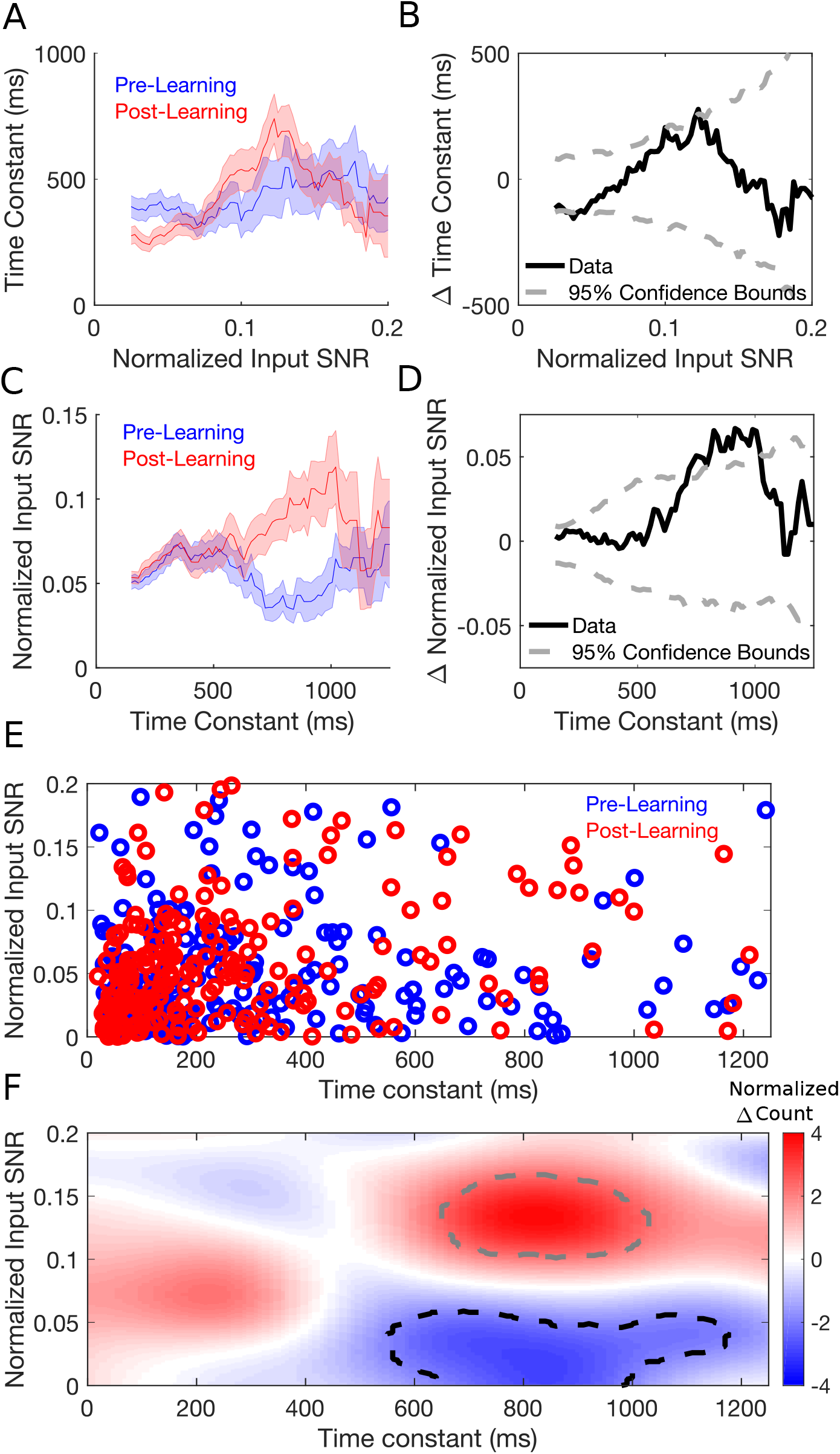
The MVAR model supports the dynamical realignment hypothesis but not the dynamical slowing hypothesis. A: Dependence of the time constants of modes on their input SNR, pre- and post-learning (average time constant conditioned on normalized input SNR, mean±sem taken over pooled modes over animals). B: Difference between pre and post curves in A (solid black line). Dashed gray lines show 2.5% and 97.5% of shuffled distributions. C, D: As in A, B but for an average of normalized input SNR conditioned on time constant. E: Time constants and normalized input SNRs of modes pooled over animals pre- and post-learning. F: Smoothed histogram of difference over learning in number of modes with a given input SNR and time constant (normalized by standard deviation over shuffles). Dashed black and gray lines show regions where the number fell below 2.5% and above 97.5% of shuffled distributions respectively (see Methods).

In principle, enhanced integration could also arise through greater non-normality in the recurrent dynamics (Supplementary Figure 1). However, we found that for 6/8 animals the recurrent dynamics became less non-normal over learning (p=0.03, two-sided Wilcoxon rank sum test), suggesting that this mechanism did not contribute to the enhancements detected in the MVAR model (Supplementary Fig 3C).

In summary, these results support the hypothesis that learning reorganizes local network interactions in order to align slowly decaying modes of recurrent dynamics with the optimal linear discriminant of sensory input (Figure 4C), thereby enhancing temporal integration of task-relevant sensory information.

### Stimulus-specific but not uniform connectivity changes reproduce the changes in dynamical integration observed in the MVAR model

How might the dynamical realignment observed in the MVAR model relate to systematic changes in synaptic connectivity and response tuning within the V1 circuit? Constraints in the original experiment meant that we were unable to determine the orientation tuning of the imaged neurons. Thus, we turned to a canonical circuit model for feature selectivity to investigate the relationship between network connectivity, tuning curves, and dynamical modes (Ben-Yishai et al., 1995; Rubin et al, 2015; Hennequin et al., 2018). The model comprised excitatory and inhibitory neurons arranged on a ring corresponding to their preferred orientation before learning. Neurons at nearby locations formed stronger synaptic connections and received more similarly tuned feedforward input than those more separated around the ring (Figure 6A). This is consistent with local microcircuits in visual cortex in which neurons receive feature-tuned feedforward input (Lien et al., 2013) and interact through feature-specific local synapses (Cossell et al., 2015; Znamenskiy et al., 2018).

**Figure 6.**
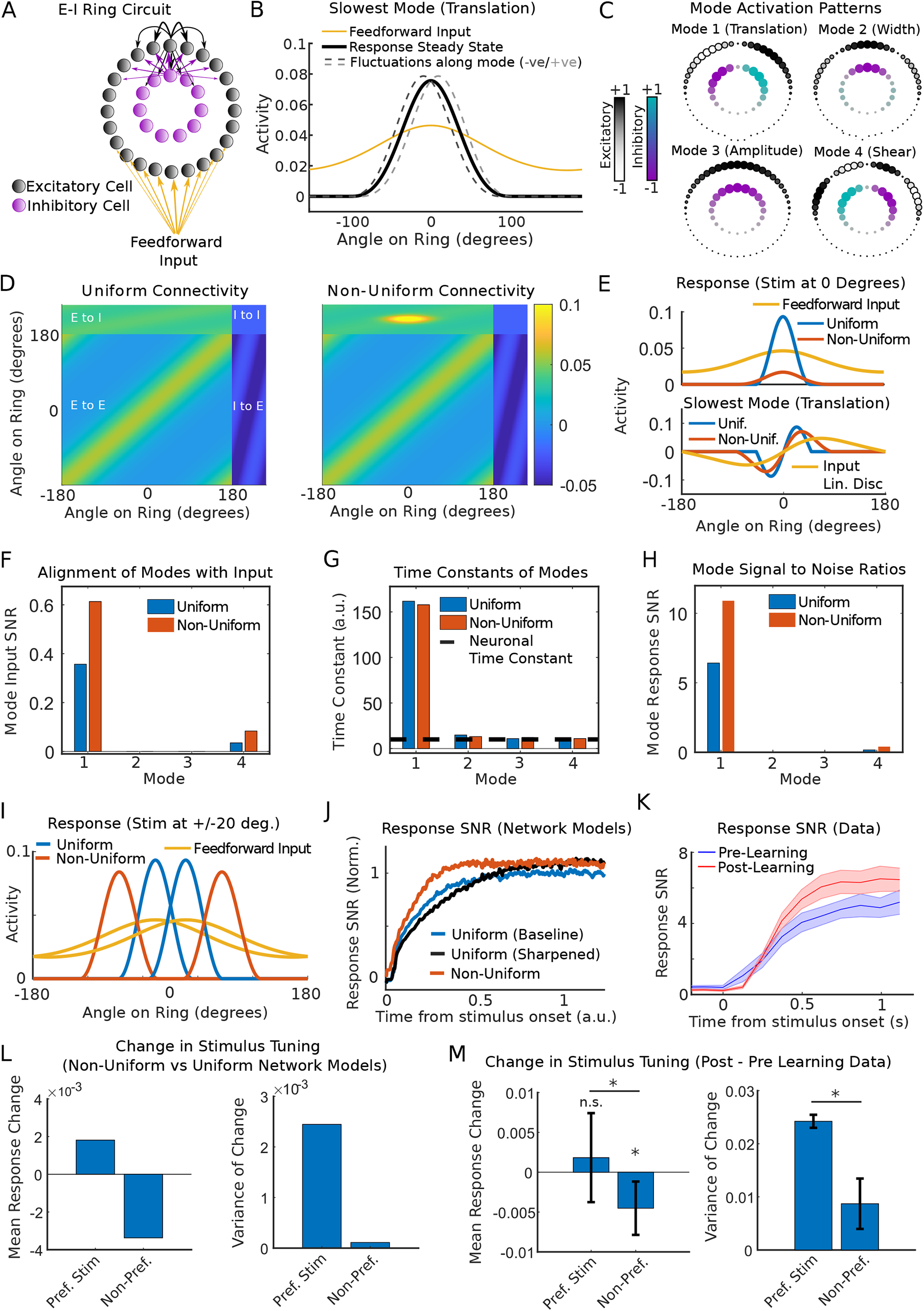
Stimulus-specific inhibition aligns the slowest decaying mode with the input linear discriminant and predicts observed changes in stimulus tuning. A: Excitatory-inhibitory ring network model for V1 orientation selectivity. B: Steady state network response (solid black) and perturbations along the most slowly decaying mode (dashed gray). Feedforward input (yellow) was rescaled for aid of visual comparison. Only excitatory cells are shown. C: Activation patterns m for the four most slowly decaying modes (in order of time constant). Size and color of circles depicts weighting of cell in mode activation pattern. D: Synaptic weight matrix for a ring network with uniform (left) and non-uniform (right) connectivity. E: (Top) Feedforward input and steady state responses for the two networks. (Bottom) The most slowly decaying mode m for each of the two networks, overlaid with the input linear discriminant. The greater overlap between red and yellow lines compared to cyan and yellow indicates increased alignment. F-J: Input SNRs (F), time constants (G) and response SNRs (H) for the four most slowly decaying modes. I: Network responses to two stimulus orientations separated by 40 degrees. J: SNR of instantaneous network output for three networks (based on simulation of nonlinear dynamics). K: SNR of imaged V1 population responses (mean±sem over mice). L: The change in responses of excitatory neurons to their preferred and non-preferred stimuli induced by non-uniform inhibition (mean and variance over cells). The greater variance for the preferred stimulus reflects a more heterogeneous response change including both boosting and suppression. M: Mean (left) and variance (right) of the change in pyramidal responses to their preferred and non-preferred stimuli over learning. Responses to the non-preferred stimulus decreased (p=0.003, two-sided sign test) but responses to the preferred stimulus did not (p=0.8, two-sided sign test; p=0.025, one-sided Wilcoxon rank sum test on difference between preferred and non-preferred stimulus response change). The variance over cells of response changes was higher for the preferred than non-preferred stimulus (p=0.035, shuffling test).

We first analyzed the tuning curves and modes of dynamics in the E-I ring network. The network formed a stable bump of activity centered on the stimulus orientation (Figure 6B, solid black line), and each of the four most slowly decaying modes reflected an interpretable fluctuation about this stable activity pattern: side-to-side translation (Figure 6B, dashed gray lines), sharpening/broadening, gain of amplitude, and asymmetric shear (Figure 6C, Supplementary Figure 4A-C). Responses were sharpened relative to feedforward input (Figure 6B, black vs yellow line) and the degree of sharpening depended on the strength and tuning of excitatory and inhibitory synapses around the ring (Supplementary Figure 4D-F). This suggested that a possible mechanism for the reorganization of dynamical modes observed in the MVAR model may be increased sharpening of feedforward input due to changes in recurrent synapses. On testing this hypothesis, however, we found that recurrent sharpening reduced alignment of the slowest dynamical mode with the input linear discriminant, in contrast to the increased alignment observed in the MVAR model (Supplementary Figure 4G-L). These findings remained consistent for a broad range of networks with varying strength and feature-tuning of synaptic weights (Supplementary Figure 5A-H). Thus, uniform changes in the strength or tuning of excitatory-excitatory and excitatory-inhibitory weights did not reproduce the changes over learning observed in the data.

We previously found that response SNRs of both excitatory and inhibitory cells increase over learning, and that these improvements are driven by an emergence of stimulus-specific excitatory to inhibitory interaction weights in the MVAR model such that E to I interaction weights amongst cells with the same stimulus preference are stronger after learning than before (Khan et al., 2018). We therefore reasoned that a change in E-I connectivity that is specific to the learned stimuli might account for the realignment of modes observed in the MVAR model. Thus, we considered a non-uniform ring network in which excitatory to inhibitory synaptic weights were strengthened locally amongst neurons tuned to a particular orientation (Figure 6D). We found that the resulting non-uniform inhibition induced changes in dynamical modes that were consistent with those observed over learning in the MVAR model: the slowest-decaying mode became better-aligned with the input discriminant while its time constant was unchanged (Figure 6E, F, Supplementary Figures 5I-L, 6A). When stimuli were presented at ±20 degrees relative to the subnetwork center (reflecting the 40-degree stimulus separation in the experiment), information was enhanced via a greater separation of responses around the ring (Figure 6I, Supplementary Figure 6B). In simulations of the full nonlinear network response to feedforward input, accumulation of stimulus information was accelerated by non-uniform inhibition but slowed by uniform sharpening (Figure 6J). Experimental data showed an accelerated rate of integration over learning consistent with the non-uniform connectivity change (Figure 6K). Thus, in both the analysis of local linearized modes and the evolution of the nonlinear network responses over time, non-uniform changes in E-I connectivity accounted for the learning-related changes in responses imaged from the V1 circuit.

The tuning curves induced by non-uniform connectivity (Figure 6I) generated further predictions that we subsequently tested on the experimental data. Responses of excitatory neurons to their non-preferred stimulus were consistently suppressed by non-uniform inhibition, whereas responses to their preferred stimulus showed a heterogeneous combination of boosting and suppression (Figure 6L). Changes over learning in imaged pyramidal cell responses showed a similar pattern (Figure 6M). Moreover, the average response SNR of both excitatory and inhibitory neurons increased in the model (Supplemental Figure 6C-F), as previously reported for the imaged responses of pyramidal cells and parvalbumin-expressing interneurons (Khan et al., 2018; reproduced in Supplementary Figure 6G).

Taken together, these findings demonstrate that the learning-related changes in imaged network responses are consistent with the emergence of stimulus-specific excitatory to inhibitory synaptic connectivity within local V1 microcircuits. These connectivity changes act to increase response information by aligning slowly decaying dynamical modes with the optimal discriminant of sensory input in order to selectively integrate relevant sensory information over time.

## Discussion

We have developed a general framework for modeling the integration and transmission of sensory information through recurrent networks and leveraged this framework to uncover the changes in recurrent processing that drive improvements in sensory representations over learning. Previous studies suggested that recurrent synapses selectively amplify or sharpen the tuning of feedforward input (Douglas et al., 1995; Ben-Yishai, 1995; Somers et al., 1995; Murphy and Miller, 2009; Liu et al., 2011; Li et al., 2013; Lien et al., 2013; Cossell et al., 2015), yet theoretical analyses concluded that sharpening reduces population response information (Seriès et al., 2004; Beck et al., 2011). Others proposed that recurrent synapses selectively suppress responses to remove redundancy between similarly tuned neurons (Olshausen and Field, 1996; Lochmann et al., 2011; Znamenskiy et al., 2018; Chettih and Harvey, 2019), yet such mechanisms cannot explain the improvements in response information as animals learn to discriminate simple sensory features such as oriented grating stimuli (Poort et al., 2015; Khan et al., 2018). Instead, we show that recurrent dynamics in primary visual cortex perform selective temporal integration of relevant sensory information, an operation previously reported only in higher sensory and non-sensory areas with longer cellular and network time constants (Shadlen and Newsome 2001; Wong and Wang, 2006; Kiebel et al., 2008; Goldman et al., 2009a; Mante et al., 2013; Murray et al., 2014).

Responses of cells in primary visual cortex have been found to decay within a single neuronal time constant when thalamic input is removed (Reinhold et al., 2015). Can the long timescales of recurrent dynamics required for selective temporal integration be reconciled with these observations? One possibility is that the dynamical regime of cortex is dependent on tonic thalamic input, or on thalamocortical loops. Alternatively, Reinhold and colleagues may have predominantly activated and measured rapidly decaying modes of dynamics which obscured the presence of slowly decaying modes intermixed with the population response. Detecting such slowly decaying modes of dynamics requires recording from neural populations, whereas Reinhold and colleagues recorded single neurons. Future studies could test these hypotheses by measuring and perturbing different patterns of population activity during sensory stimulation and quantifying the time constants of network responses.

We inferred cortical dynamics by fitting linear dynamical models to imaged population activity. Such an approach is prone to model mismatch, such that temporally coordinated external input may be erroneously attributed to local interactions amongst cells. Thus, while we identified changes in dynamics over learning, it is possible that such dynamics are inherited by the local circuit or generated through a broader network of cortical and subcortical structures. This hypothesis could be tested in future experiments by recording neuronal population activity in multiple brain regions simultaneously during sensorimotor decision-making tasks. Additional confounds may arise through the convolution of neuronal responses by slow calcium dynamics and the temporal resolution of the data (∼125 ms). However, although these may lead to an overestimate of the time constants of network dynamics, they cannot trivially explain the change in alignment of dynamical modes observed over learning. Nonetheless, while we observed an apparent decrease in non-normality over learning, measurements at higher temporal resolution are necessary to detect rapid forms of non-normal dynamics and their changes over learning (Murphy and Miller, 2009).

Our theory explains a recent report that information-limiting noise correlations are higher when animals make correct decisions compared to incorrect ones (Valente et al., 2021). Because these correlations reduce the information about the stimulus available in the network response relative to an uncorrelated population and yet were associated with improved behavioral accuracy, these findings were considered to be paradoxical by Valente and colleagues. Instead, we show that these findings are an expected signature of optimal integration of sensory input through the recurrent circuit dynamics. In particular, we observe that information-limiting response correlations across neurons are maximized when networks integrate their sensory input optimally (compare Figure 1F to Figure 1H and Supplementary Figure 1A, ellipses which are more elongated along the direction which separates the two means have higher information-limiting correlations). Valente and colleagues also found that correlations between responses at different time points within a trial are higher when animals make correct decisions, which was considered paradoxical because such correlations limit the ability of downstream readers to decode the stimulus over the duration of a trial. We show that strong temporal correlations are an expected signature of optimal integration of sensory input through time by the circuit. Thus, we suggest that optimal sensory coding is best understood in terms of the transformation of sensory input signals by the neural circuit, a perspective which leads to fundamentally different experimental predictions for the optimal response statistics than those obtained using abstract neural encoding models (see also Seriès et al., 2004; Beck et al., 2011; Huang et al., 2020).

Several previous studies have investigated information transmission through recurrent networks (Seriès et al., 2004; Ganguli et al., 2008; Beck et al., 2011; Toyoizumi and Abbott, 2011; Dambre et al., 2012; Najafi et al., 2018; Huang et al., 2020). While most studies (correctly) concluded that information in network output cannot exceed that contained in the input, such studies either 1) quantified information in time-integrated network responses (Seriès et al., 2004; Moreno-Bote et al., 2014), 2) modeled sensory input as being static within each trial, varying only from trial to trial (Najafi et al., 2018), or 3) analyzed network models which lack the capacity for dynamical integration (Beck et al., 2011). In our analysis, input noise was time-varying and recurrent dynamics could integrate input over the course of a trial, allowing the instantaneous (but not time-integrated) response information to exceed that of the input. While Toyoizumi and Abbott considered a similar scenario, their analysis was restricted to networks of randomly connected neurons with antisymmetric, saturating transfer functions.

Our analysis provides a general framework for understanding evidence integration in neural circuits, such as path integration in grid cells, vestibular integration in head direction cells, and integration of motion in higher visual areas. While several of these systems have been studied mechanistically as attractor networks (Wong and Wang 2006; Burak and Fiete, 2009) or statistically as drift-diffusion and population coding models (Ratcliff and McKoon, 2008; Averbeck et al., 2006), our approach provides a unifying formalism which links statistical properties of evidence integration and population coding to the dynamical properties of the underlying recurrent network. While we have focused on changes in network dynamics over learning, the mechanism of dynamical alignment may also provide a substrate for contextual or attentional modulation of sensory processing (Gilbert and Li, 2013). Specifically, top-down input may modulate the dynamics of recipient neural populations, transiently aligning dynamical modes of the local circuit with relevant features of bottom-up sensory input according to task context. Such a mechanism could allow for flexible routing and gating of information between brain areas through the dynamical formation and coordination of “communication subspaces” (Semedo et al., 2019; Kohn et al., 2020; Javadzadeh and Hofer, 2021), configured through selective alignment of local modes across anatomically distributed circuits.

## Methods

### Resource Availability

#### Lead Contact

Further information and requests for resources and reagents should be directed to and will be fulfilled by the lead contact and corresponding authors Angus Chadwick and Maneesh Sahani.

#### Materials Availability

This study did not generate new unique reagents

#### Data and Code Availability

The data and code that support the findings of this study are available from the corresponding authors upon request.

### Experimental model and subject details

No new experimental data were collected for the purposes of this study. The acquisition and pre-processing of data used in this study are described in detail in Khan et al., 2018.

### Method details

#### Analysis of optimal stimulus discrimination function (Figure 1)

In the Supplementary Mathematical Note we analyze the problem of stimulus discrimination from a signal processing perspective. We consider a network receiving noisy but stimulus-tuned input and tasked with reporting stimulus identity in its output. Under the assumption that the input time series for a given stimulus follows a multivariate normal distribution with temporally uncorrelated, stimulus-independent noise, we show that the statistically optimal method for discriminating two stimuli is to perform a linear projection and temporal filtering of the input time series. We derive the optimal projection weights and filter, and the signal to noise ratio (SNR) obtained using an arbitrary projection and filter.

In Figure 1 we sought to illustrate these observations in a minimal toy example consisting of a reduced two-dimensional system describing the feedforward input to two neurons under each of two stimuli. The dimensionality and statistics of the input were chosen primarily to optimize visualisation and conceptual insight - our analysis allows for arbitrary numbers of neurons receiving input with arbitrary stimulus-tuning and noise covariance. For each stimulus *s*_*i*_ (*i* = 1, 2) and at each timestep *t*, feedforward inputs **u**(*s*_*i*_, *t*) ∼ *N*(**g**(*s*_*i*_), Σ_*η*_) were sampled independently from a multivariate normal distribution with stimulus-dependent mean **g**(*s*_1_) = [1, 2], **g**(*s*_2_) = [2, 1] and stimulus-independent covariance Σ_*η*_ = [0, 2; 2, 1] (here and throughout, we will use the shorthand notation that matrix elements separated by commas are on the same row, while elements separated by a semicolon are on separate rows, e.g. [*x, y*] = [*x*; *y*]^*T*^). These time series were projected onto the linear discriminant 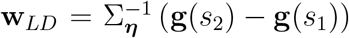 to obtain 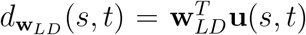 before being summed cumulatively over time to obtain 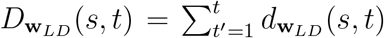. The signal (difference in mean), noise (standard deviation), and signal to noise ratio of the projection of instantaneous input onto a vector **w**, *d*_**w**_(*s, t*) = **w**^*T*^ **u**(*s, t*), were plotted using analytical expressions Δ*μ*_input_(**w**) ≡ ⟨*d*_**w**_(*s*_2_, *t*) − *d*_**w**_(*s*_1_, *t*)⟩ = **w**^*T*^ (**g**(*s*_2_)−**g**(*s*_1_)), 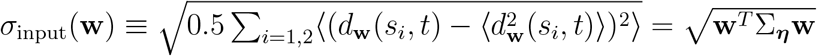, SNR_input_(**w**) = Δ*μ*_input_(**w**)/*σ*_input_(**w**). Following temporal integration, the corresponding quantities 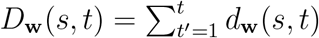 were plotted as Δ*μ*_input_(**w**, t) ≡ ⟨*D***w**(*s*2, *t*) − *D***w**(*s2, t*)⟩ = Δ*μ*_input_(**w**)*t*, 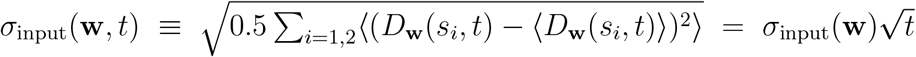 and 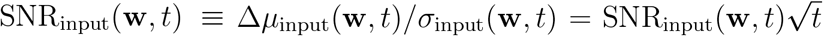. Iso-probability contours at one standard deviation under each stimulus were plotted as 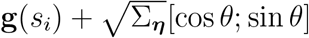

#### Analysis of linear Fisher Information in recurrent networks (Figure 2 and Supplementary Figure 1)

Linear Fisher Information quantifies the accuracy of a locally optimal linear estimator of a stimulus from network responses (Seriès et al., 2004; Beck et al., 2011). When network responses follow a multivariate normal distribution, the linear Fisher Information takes the form of a (squared) signal to noise ratio. We derived analytical expressions for the linear Fisher Information of the instantaneous output of a recurrent network as a function of its input statistics and dynamics, and for the SNR of network output projected onto any one its dynamical modes (see Supplementary Mathematical Note). Our results hold for networks with arbitrary numbers of neurons with arbitrary nonlinearities and synaptic connectivity, receiving sensory input with arbitrary stimulus-tuning and noise covariance. Our strongest modeling assumptions were the linearization of dynamics about a fixed point and the analysis of stationary state response statistics.

##### Signal to noise ratio along dynamical modes (Figure 2)

To illustrate the relationship between network dynamics and population coding, we constructed a minimal toy model comprising a two-dimensional linear dynamical system 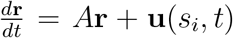 corresponding to the linearized dynamics of the firing rates **r** = [*r*_1_; *r*_2_] of two reciprocally connected neurons. The weight matrix *A* was constructed by defining two dynamical modes with activation patterns **m**_*i*_ and corresponding time constants *τ*_*i*_. We consider a system without oscillations, i.e. one in which the eigenvalues *λ*_*i*_ of *A* are real. In that case, *τ*_*i*_ = −1/*λ*_*i*_ and the unique weight matrix which generates these dynamical modes is given by *A* = *M*^−1^Λ*M*, where 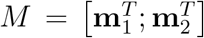 and Λ = [*λ*_1_, 0; 0, *λ*_2_] (note that we define the mode activation patterns **m**_*i*_ to be the *left eigenvectors* of *A*, see Supplementary Mathematical Note for details). We constructed **m**_*i*_ as unit length vectors with a given angle relative to the input linear discriminant using the equation **m**_*i*_ = *R*(*θ*_*i*_)**w**_*LD*_/‖**w**_*LD*_‖, where *R*(*θ*_*i*_) = [cos(*θ*_*i*_), −sin(*θ*_*i*_); sin(*θ*_*i*_), cos(*θ*_*i*_)] is a rotation matrix. **w**_*LD*_ was defined as the linear discriminant of two stimulus inputs with **g**(*s*_1_) = [6; 6], **g**(*s*_2_) = [5; 7], Σ_*η*_ = [20, 10; 10, 20] (these values, along with the modes and time constants, were chosen to primarily to optimize visualisation). We constructed networks with one mode aligned to input linear discriminant and the other orthogonal to the first by setting *θ*_1_ = 0.02*π, θ*_2_ = *θ*_1_ + 3*π*/2. For the network with slowly-decaying mode aligned to the linear discriminant we set *τ*_1_ = 10, *τ*_2_ = 2, and for the network with rapidly-decaying mode aligned to input linear discriminant we set *τ*_1_ = 2, *τ*_2_ = 10 (in arbitrary units of time).

As panels A-C were designed to illustrate the dynamical modes of the network rather than the stimulus input, we set the input to **u** = (**g**(*s*_1_) + **g**(*s*_2_))/2 (or **u** = [0; 0] before input onset). Network responses **r** were computed using the solution to the linear dynamics **r**(*t*) = exp(*A t*)(**r**(0) − **r**_∞_) + **r**_∞_ where **r**(0) = [0; 0], **r**_∞_ = −*A*^−1^**u** and exp is the matrix exponential function. The perturbation was modeled by setting **r**(*t*_pert_) = **r**_∞_ + [0; 10] and computing all future time points as **r**(*t*) = exp(*A*(*t* − *t*_pert_))(**r**(*t*_pert_) − **r**_∞_) + **r**_∞_

For panels D-J, network responses to the two stimulus input time series were simulated using the Euler method with *dt* = 0.01, i.e. **r**(*t* + *dt*) = **r**(*t*) + (*A***r**(*t*) + **g**(*s*_*i*_) + *η*_*t*_)*dt* where *η*_*t*_ ∼ *N*(0, Σ_*η*_). For visualisation purposes, trajectories were smoothed before plotting for panels E and G using a moving average box filter containing 100 time samples.

Input and output iso-probability ellipses were generated as in Figure 1, using the relevant mean and covariance matrix in each condition. Response means were computed using the analytical solution for a linear system at steady state, **r**_∞_(*s*) = −*A*^−1^**g**(*s*), and response covariance matrices (panels F and H) were computed as the solution to the Lyapunov equation *A*Σ + Σ*A*^*T*^ + Σ_*η*_ = 0 using the Matlab function *lyap*.

The signal, noise, and signal to noise ratio of stationary state responses projected along each mode 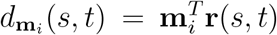 were computed using the equations Δ*μ*_output_(**m**_*i*_) ≡ ⟨*d*_**m**_ (*s*_2_, *t*) − *d*_**m***i*_ (*s*_1_, *t*)⟩ = Δ*μ*_input_(**m**_*i*_)*τ*_*i*_, 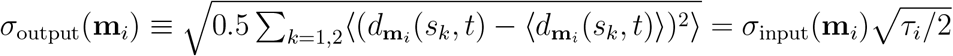 and 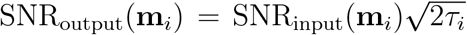 respectively, where Δ*μ*_input_, *σ*_input_, SNR_input_ are as described for Figure 1 (see Supplementary Mathematical Note for a derivation).

##### Non-normal dynamics (Supplementary Figure 1)

We derived expressions relating linear Fisher Information to the dynamics of an arbitrary normal or non-normal network (subject to the same approximations described above). These expressions had a simple and interpretable form in three special cases: two-dimensional networks, normal networks, and non-normal networks with strong functionally-feedforward dynamics. Related findings have been presented previously (Ganguli et al., 2008; Goldman et al., 2009).

To illustrate our analytical findings for the two-dimensional case, we constructed networks with modes **m**_1_ = [cos *θ*_1_; sin *θ*_1_], **m**_2_ = [cos *θ*_2_; sin *θ*_2_]. Panel A was constructed using the same procedure as for Figure 2, but this time with *τ*_1_ = 10, *τ*_2_ = 5. For panel B we chose input with isotropic covariance Σ_*η*_ = ***I***_2_ (where ***I***_*N*_ is the N x N identity matrix) and Δ**g** = **g**(*s*_2_) − **g**(*s*_1_) = [1; 0]. These inputs were chosen in order to demonstrate the influence of non-normality as clearly as possible. We set *τ*_1_ = 10, *τ*_2_ = 1, 5, 7.5, 9 and varied *θ*_1_, *θ*_2_ from 0 to *π* for each value. For each network (defined by the parameters *θ*_1_, *θ*_2_, *τ*_1_, *τ*_2_ using the procedure described for Figure 2), the Fisher Information of the stationary state network response *ℐ*_*F*_ = Δ**r** · Σ^−1^Δ**r** was computed by substituting the long-run solution for the mean Δ**r** = −*A*^−1^Δ**g** and the numerical solution to the Lyapunov equation for Σ (described above). We normalized this linear Fisher Information by the maximum achievable SNR in any normal network with the same time constants by defining 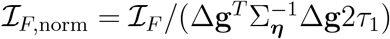.

To illustrate the case of functionally-feedforward networks (Goldman et al., 2009), we constructed networks with NxN weight matrix *A*_*ij*_ = (−1/*τ*)*δ*_*ij*_ + *ωδ*_*i,j*+1_, while varying the weight *ω* and number of neurons *N* for fixed single-cell time constants *τ* = 10 (where *δ*_*ij*_ is the Kronecker delta symbol). We set Δ*g*_*i*_ = *δ*_*i*1_ and Σ_*η*_ = ***I***_*N*_. We derived analytical expressions in the *ω* → ∞ limit for the linear Fisher Information of network output at stationary state, the temporal filter the network applies to its input, and the optimal linear readout of network responses. We numerically extended our results to the finite *ω* case by computing the response signal, response covariance, and linear Fisher Information in the same way as for the two-dimensional networks. To understand how the finite *ω* and large *ω* networks differ and where the large *ω* approximation breaks down, we also computed the SNR of the finite *ω* network responses projected onto the large *ω* optimal readout. Full derivations can be found in the Supplementary Mathematical Note.

#### Multivariate autoregressive system model and analysis of neural data (Figure 3, 5, Supplementary Figure 2, 3)

Details of the experiment, data preprocessing, calculation of behavioral d-prime (Figure 3B), and fitting and validation of MVAR model on this dataset data have been described in detail in previous publications (Khan et al., 2018; see also Poort et al., 2015, 2021). Here, we summarize the MVAR model and provide details of novel MVAR analyses used in the present study.

The imaged Δ*F*/*F* signals for each cell were divided into trials of duration -1 to 1 s relative to the onset of a visual stimulus. Here and below, all sums over time samples are restricted to the *N*_t_ = 9 time samples contained in the post-stimulus window of 0 to 1 s (although the model was fit to the full window of -1 to 1 s containing 17 time samples). We collect the population activity of *N* simultaneously imaged neurons at imaging frame *t* on trial *i* into an *N*-dimensional vector denoted 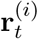. We define the following quantities which we will make use of below. The trial-averaged activity conditioned on stimulus *S* and time relative to stimulus onset *t* is 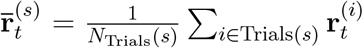, where *N*_Trials_(*s*) is the number trials of stimulus *S*. The grand average over both time samples and trials conditioned on the stimulus *s* is 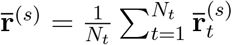. The pooled covariance over vertical (*V*) and angled (*A*) stimuli is 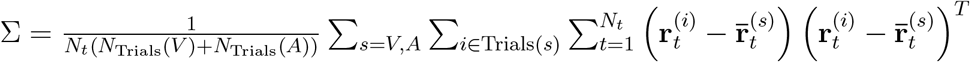.

##### Description of Model

To infer linear dynamics and stimulus input of the imaged circuit, we fit a multivariate autoregressive linear dynamical system model to the imaged responses. In the MVAR model, the imaged activity is modeled as:

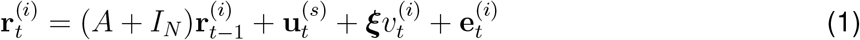

where *A* is an *N* × *N* matrix of interaction weights, 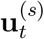 is a vector of *N* stimulus-related inputs, *ξ* is a vector of *N* running speed coefficients, 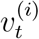 is the running speed of the animal and 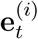 is a vector of residuals.

The MVAR model is fit to each dataset by minimizing the sum of squared residuals across all neurons and trials of the vertical, angled, and gray corridor stimuli before or after learning (−1 to 1 s about the onset of the corridor, which appeared suddenly). Analytical expressions for the model parameters obtained under this least squares fit offer insight into their interpretation (equations 2-4 in Khan et al., 2018). In particular, the interaction weights depend only on the stimulus-independent covariance of the data (both the instantaneous covariance Σ and the covariance between consecutive imaging frames). Given these interaction weights, the stimulus-related input depends only on the stimulus-conditioned trial-averaged responses 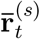. Thus, the MVAR model uses the imaged noise covariance of the data (both within and across consecutive time samples) in order to infer interactions between cells, and ascribes any remaining stimulus-dependent variation in trial-averaged responses to sensory input. The residuals have zero mean under each condition, i.e. 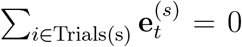 for any and*s* (equation 4 in Khan et al., 2018). We observed that the contribution of the running speed term to responses was negligible and so do not report results on this term (note that *ξ* was constrained to have the same value pre- and post-learning in all of our analyses - when *ξ* was free to vary over learning a larger contribution could be observed).

##### Visualization of MVAR input and output along discriminant axis

Having fit the MVAR model to the experimental data, we sought to visualize how the imaged responses were generated through recurrent integration of stimulus-related input within the inferred dynamical system. To do so, we projected the sensory input, recurrent input, and MVAR output onto the linear discriminant in order to see how stimulus-discriminability evolved over time. Single-trial sensory input was defined as 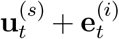 (i.e. residuals were assigned as input noise), recurrent input as 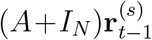, and MVAR output as 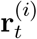. The linear discriminant vectors were 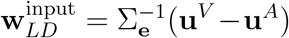 and 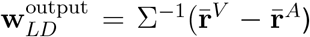, where 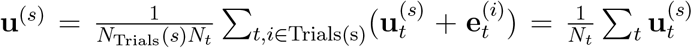 and 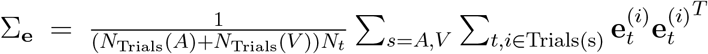. The sensory input was projected onto 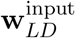, while both recurrent input and imaged responses were projected onto 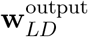. We plotted the mean and standard deviation over trials of these projected activity patterns for a representative mouse in the post-learning condition.

##### Quantification of MVAR input and output information

The stimulus-information (or linear discriminability) of single-imaging frame population responses was quantified as 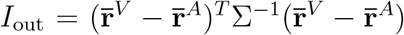. The stimulus-information of inferred input was quantified as 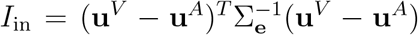. These metrics were computed separately for the pre- and post-learning data for each mouse. The gain in output to input information was defined as 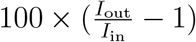.

##### Quantification of temporal integration of relevant and irrelevant input

To test how temporal integration of relevant and irrelevant input changed over learning in the MVAR model, we analyzed the impulse-response of the MVAR to two different input perturbations. The impulse-response to a perturbation **p** was modelled by setting the MVAR to an initial state **r**_0_ = **p** and forward-simulating the system over multiple time steps with no other input, i.e. **u**_*t*_, **e**_*t*_, *v*_*t*_ = 0. This gave the response **r**_*t*_ = (*A* + ***I***_*N*_)^*t*^**p**. Simulated responses **r**_*t*_ were then projected onto a vector **w**. For the relevant input, we chose **p** to be the MVAR input linear discriminant 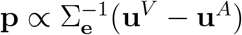 and **w** to be the linear discriminant of the imaged population responses 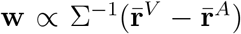. With this choice (i.e., by choosing not to enforce **w** = **p**), we allow for the possibility that temporal integration occurs through either normal or non-normal dynamics (Supplementary Figure 1). For the task-irrelevant input we chose 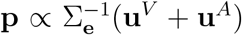 and 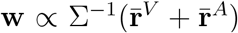. Time constants of network responses were defined as 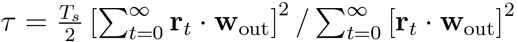, which was adapted from the analytically-derived temporal integration factor ***I***_*T*_ (*f*) in the Supplementary Mathematical Note (see section titled Signal Processing Analysis).

##### Constrained model fits

To test whether the learning-related changes in temporal integration in the MVAR model require changes in interaction weights or stimulus input, we refit the MVAR with either *A* or **u** constrained be the same both pre- and post-learning. We then repeated the analyses for Figure 3 on the constrained MVAR model fits. Details of the constrained model fitting procedure are provided in Khan et al., (2018).

##### Input and output SNR along MVAR modes

To compute the SNR of network input and output projected onto each mode, we used analytically derived expressions which relate these SNRs to the eigenvectors and eigenvalues of *A*. Eigenvectors (right 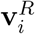 and left 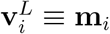) and eigenvalues *λ*_*i*_ of the pre- and post-learning MVAR interaction weight matrices *A* were numerically computed using the Matlab function *eig*. The SNR of stimulus input projected along each mode was then given by the equation 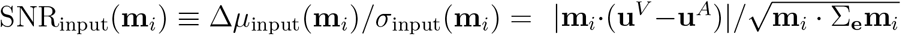. The normalized input SNR was SNR_norm_(**m**_*i*_) = SNR_input_(**m**_*i*_)/SNR_input_(**w**_*LD*,input_) where 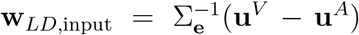 is the input linear discriminant and 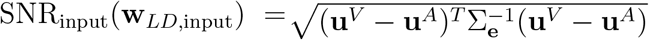 is the SNR of input projected along the linear discriminant. We computed the time constant of each mode using the equation *τ*_*i*_ = − *T*_*s*_/ log (*λ*_*i*_ + 1) which converts from a discrete-time dynamical system of sampling period *T*_*s*_ to a time constant in an equivalent continuous-time dynamical system. We restricted our analysis of individual modes to those with real eigenvalues *λ*_*i*_ + 1 > 0 (which ensures that *τ*_*i*_ are real, so that the mode is not oscillatory).

We pooled modes across animals separately in the pre- and post-learning conditions (note that individual modes are not matched pre- vs post-learning). Both pre- and post-learning, we performed averages over time constants conditioned on normalized input SNRs and over normalized input SNRs conditioned on time constants. These conditional averages were obtained using a moving average analysis. To obtain an average normalized input SNR conditioned on time constant, we used a box filter of width 100 ms with center increasing from 100 ms to 1400 ms in increments of 25 ms. For each increment, we computed the mean normalized input SNR of all modes within that window. Similarly, we used a box filter of width 0.025 increasing from 0.025 to 0.25 to compute average time constant conditioned on normalized input SNR. As an additional analysis, we computed a two-dimensional histogram describing the number of modes *n*(*τ*, SNR_norm_) with time constant *τ* and normalized input SNR SNR_norm_ by applying a moving two-dimensional Gaussian filter over the set of modes using the equation 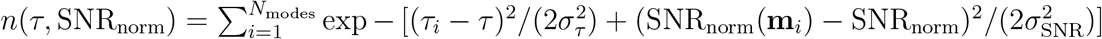. We set *σ*_*τ*_ = 100 ms and *σ*_SNR_ = 0.025. We computed the change over learning Δ*n* = *n*_post_ *n*_pre_ and normalized this quantity by its standard deviation across shuffled data (see below) to obtain Δ*n*/*σ*(Δ*n*_shuff_), a measure of the change relative to chance level, which is plotted in Figure 5F.

To determine whether learning-related changes in time constants or normalized input SNRs exceeded chance level, we performed a bootstrapping procedure based on shuffling of trials. For each mouse, we pooled pre- and post-learning trials and randomly resampled (without replacement) two sets of trials of equal number to the pre- and post-learning datasets. These shuffled datasets constituted the null hypothesis that no changes occurred over learning. We then refit the MVAR model to each of these shuffled datasets and repeated the above analyses to obtain the time constants and normalized input SNRs under the null hypothesis. In this way, we generated a null distribution for each statistic (moving average of change in time constant, moving average of change in normalized input SNR, and Δ*n*). We then formed 95% confidence intervals for each statistic based on their respective null distributions. Our null distributions consisted of 1000 such shuffles.

To confirm that our results were not biased by individual mice, we also performed within-animal averages of the time constants and normalized input SNRs pre- and post-learning (Supplementary Figure 3A,B). For this analysis, individual mice rather were considered as the statistical unit when performing significance testing.

##### MVAR non-normal dynamics

The non-normality of dynamics was quantified using Henrici’s departure from normality (Henrici, 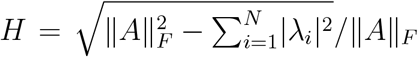, where ‖*A*‖_*F*_ is the Frobenius norm. This measure was computed separately on the interaction weight matrix for pre- and post-learning data for each animal (Supplementary Figure 3C).

#### Network models (Figure 6, Supplementary Figure 4-6)

##### Model Description

We considered two populations of cells (excitatory and inhibitory), each arranged on a ring, with *N*^*X*^ cells in population *X* ∈ {*E*, ***I***}. Each population is parameterized by its orientation on the ring 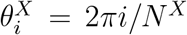. Dynamics were governed by the Wilson-Cowan equation 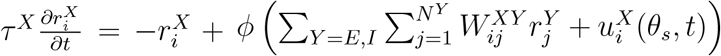, where 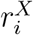 is the firing rate of neuron *i* in population *X, τ* ^*X*^ is the time constant of neurons in population 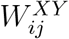 is the weight from neuron *j* in population *Y* to neuron *i* in population *X*, 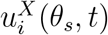 is the external input to neuron *i* in population *X* as a function of the stimulus orientation *θ*_*s*_ and time *t*, and *ϕ* is an element-wise nonlinearity. For both *E* and ***I*** populations we used a threshold-power law nonlinearity 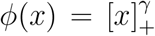 (Hansel and Van Vreeswijk, 2002; Miller and Troyer, 2002; Ahmadian et al., 2013; Rubin et al., 2013; Hennequin et al., 2018).

External input had stimulus-tuned mean 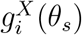 and additive, temporally uncorrelated Gaussian noise 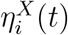, i.e. 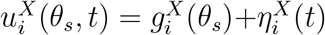 with 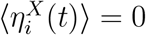 and 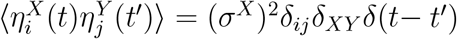. Input tuning curves were circular-Gaussian, rotationally-invariant functions of stimulus orientation, defined by von Mises functions 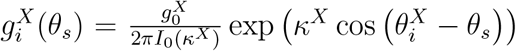. The parameter *κ*^*X*^ determines how concentrated the inputs are around the ring (i.e., orientation selectivity of input), while 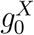 controls the total strength of network input. ***I***_0_ is the modified Bessel function of the first kind, which is included to normalize the total input strength so as to be independent of the input tuning *κ*^*X*^. To preserve rotational symmetry, inputs were chosen such that that 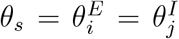 for some pair of integers *i, j*.

For the uniform network, weights had the same circular-Gaussian form as the input, 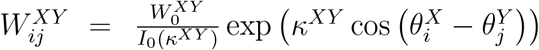 where *κ*^*XY*^, 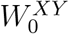 are the concentration and strength parameters for the weights from population *Y* to population *X*. For the non-uniform network, the excitatory to inhibitory weights were modified to 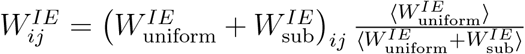 where 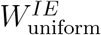 is the connectivity for the uniform network, 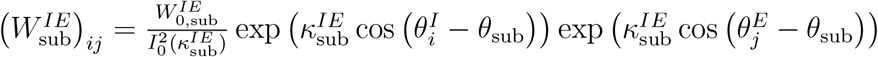 is the additional subnetwork connectivity, ⟨*W* ⟩ denotes an average over all elements of the weight matrix *W* and *κ*_sub_, 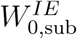 are the concentration and strength parameters for the excitatory-inhibitory subnetwork.

With the exception of parameter sweeps, all analyses of the uniform and non-uniform network used the following parameters: *N*^*E*^ = 1000, *N* ^***I***^ = 200, *τ* ^*E*^ = 10, *τ* ^***I***^ = 5, *γ* = 2, *κ*^*E*^ = 0.5, *κ*^***I***^ = 0, 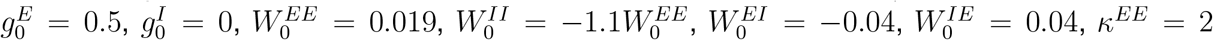, *κ*^***II***^ = 0, *κ*^*IE*^ = 0.1, *κ*^*E****I***^ = 0.4, 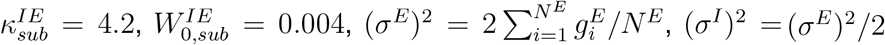. For parameter sweeps, all parameters other than those varied were held at these values. In Supplementary Figure 4, the network with weak sharpening used *κ*^*EE*^ = 1.4, *κ*^*IE*^ = 0.9, while the network with strong sharpening used *κ*^*EE*^ = 2.8, *κ*^*IE*^ = 0.4, with all other parameters unchanged.

##### Analysis of linearized dynamics

In order to compute modes of linearized dynamics and their time constants we used numerical methods to find the fixed points of the network dynamics and then numerically computed the eigenvalues and eigenvectors of an analytically-derived Jacobian.

We found that fixed point estimates obtained by forward-simulating with the Euler method yielded in-accurate estimates of linearized dynamics. Instead, we found the fixed points of Equation (4) using a root-finding algorithm applied to the equation 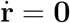, where **r** = [**r**^*E*^; **r**^***I***^], *W* = [*W* ^*EE*^, *W* ^*E****I***^; *W* ^*IE*^, *W* ^***II***^] etc., *T* is a diagonal matrix of neuronal time constants, and 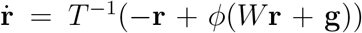. We used Newton’s method with the analytically-derived Jacobian 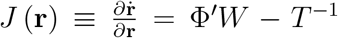 (where Φ′ = *T* ^−1^diag(*γϕ*(*W* **r** + **g**)^1−1/*γ*^) for our choice of transfer function). Fixed point estimates **r**_*n*_ were iteratively updated as 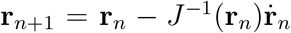. The algorithm was terminated when 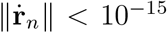 (where it was considered to have converged), or after 100 iterations (which was classed as a failure to converge). The root-finding algorithm was initialized at **r**_0_ = 0 (or when performing a parameter sweep, at the fixed point obtained from the previous set of parameters).

Having found a fixed point, the time constants, input SNRs, and output SNRs of linearized dynamical modes were computed using analytically-derived equations 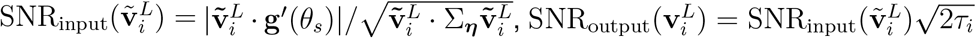, where *λ*_*i*_, 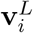, are eigenvalues and left eigenvectors of the Jacobian 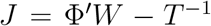, and 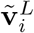 are the left eigenvectors of the matrix 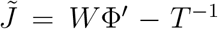. Note that 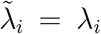, and that Φ′ = *T* ^−1^diag(*γ*r^1−1/*γ*^) at the fixed point (see Supplementary Mathematical Note). Where modes are explicitly plotted (Figures 6B, C, E, Supplementary Figure 4A-D, G-I, Supplementary Figure 6A), the quantities shown are the elements of 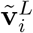. The normalized input SNR was computed as 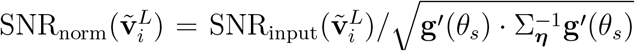. The degree of recurrent sharpening was quantified as 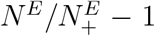, where 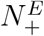 is the number of excitatory neurons with non-zero firing rate at the fixed point.

##### Analysis of two-stimulus discrimination and nonlinear dynamics

Our theoretical results are underpinned by two key approximations: the linearization of network dynamics about a fixed point and the analysis of stationary state response statistics of the linearized system. The linearization of dynamics restricts the domain of application of our theory to fine-scale sensory discrimination tasks, whereas the stimuli presented experimentally were separated by 40^°^. We therefore sought numerically determine whether our linearized theory provides adequate insight into the full nonlinear and non-stationary integration of the experimentally presented stimuli through the recurrent network. We took two approaches to do this. First, to determine the stationary state response information for two stimuli separated by 40^°^, we separately computed the linearized stationary state response statistics about each stimulus (Figure 6I and Supplementary Figure 6B-F) and then used linear discriminant analysis to compute response information. Second, to determine the non-stationary integration of input through the network dynamics following stimulus onset, we numerically computed responses of the nonlinear system over time using the Euler method (Figure 6J). The behavior of the linearized system made predictions that we were able to confirm in simulations of the nonlinear system: recurrent sharpening caused the most slowly-decaying mode to increase its time constant and become less aligned with the input discriminant (Supplementary Figure 4), which predicts that input information should be integrated more slowly but over a longer time window, and should should nonetheless achieve a greater stationary state information relative to the non-sharpened network; similarly, non-uniform inhibition caused the most slowly-decaying mode to become better aligned to the input discriminant without changing its time constant (Figure 6E-H), which predicts that input information should be integrated more rapidly, with response information reaching its plateau before the sharpened or baseline uniform network. Both predictions were borne out in simulations of the non-stationary nonlinear dynamics (Figure 6J), which demonstrates that the linearized stationary state approximation to the network dynamics is able to adequately capture the qualitative behavior of the integrative behavior of the nonlinear non-stationary system. We then verified that the same qualitative behavior could be observed in the data (Figure 6K), as would be expected based on the observed changes in MVAR modes (Figure 4).

For Figure 6I and Supplementary Figure 6B-F we computed the fixed points and Jacobians associated with the two stimulus orientations 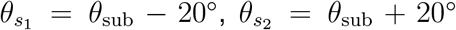. We computed stationary state response covariance around each of these fixed points by numerically solving the corresponding Lyapunov equation *J*Σ+Σ*J*^*T*^ +Φ′Σ_*η*_Φ′ = 0. We computed response information as 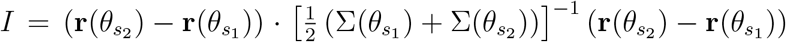. Response information was then normalized by the response information computed for a network with 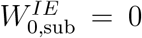 (computed using the same method with all other parameters unchanged). The SNR of excitatory and inhibitory responses were computed as 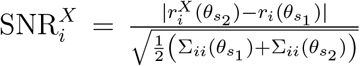. In Supplementary Figure 6C, D, we plotted 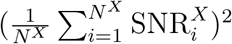 normalized by the its value in the network with 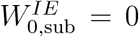 in order to facilitate direct comparison with the response information. In Supplementary Figure 6E we plotted the unnormalized 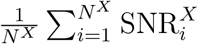 to facilitate comparison with previously defined measures of neuronal response SNR (see Khan et al., 2018, in which this measure is reported as the mean absolute selectivity).

To investigate the non-stationary and non-linear integration of sensory input following stimulus onset, we numerically solved the Wilson-Cowan equation using the Euler method. We used a time step of *dt* = 1 and initialized the simulation at the fixed point **r**(*θ*_sub_) with external input given by one of the two stimuli 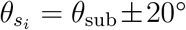. At each time step we computed the projection of responses onto the stationary state linear discriminant 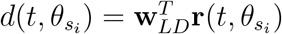, with 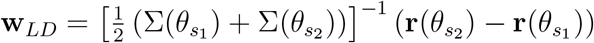 computed using the analytical equations for the stationary state means and covariances in the linearized systems about each fixed point. We simulated 1000 trials with 1000 time steps each. We computed the signal-to-noise ratio of this quantity as 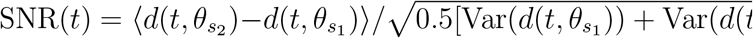 where averages and variances were taken over trials at each point in time. For the baseline and non-uniform networks we set *κ*^*EE*^ = 1.8, and for the sharpened network *κ*^*EE*^ = 2. For the non-uniform network we set 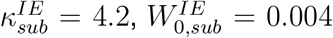 and for the baseline and sharpened network 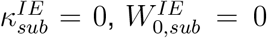. We normalized SNR(*t*) by the average value in the final 300 time steps under the baseline network model.

To compute response SNR as a function of time in the experimental data, we computed the linear discriminant as 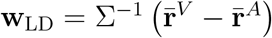 where Σ and 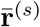 were computed as in Figure 3. We projected imaged responses 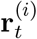 onto **w**_LD_ at each time point *t* on each trial for the vertical and angled stimuli to obtain 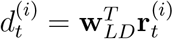. We computed the signal-to-noise ratio of this projection at each time point relative to stimulus onset by computing its mean difference between stimuli and its pooled standard deviation across stimuli, i.e. 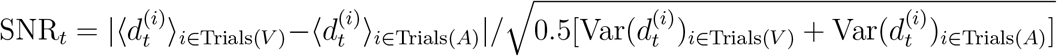. We performed this analysis separately for the pre- and post-learning data for each animal.

##### Comparison of response changes to preferred and non-preferred stimuli in model and data

We computed the change in the response of excitatory and inhibitory cells to their preferred and non-preferred stimuli over learning (in the experimental data) and between the uniform and non-uniform ring network models.

In the network models, we defined the preferred stimulus of excitatory cell *i* as the stimulus which generates the greater firing rate value at the fixed point, i.e. 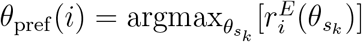 where *k* = 1, 2. The change in response to its preferred stimulus was defined as the difference in response between the two networks, i.e. 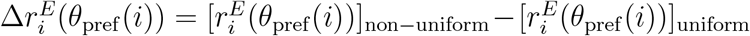 (note that cells did not change stimulus preference). The mean and variance of this change in response were then taken over the population of excitatory cells, i.e. 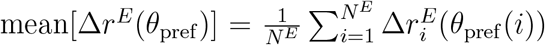, and 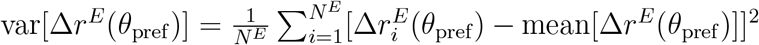. The non-preferred stimulus was analyzed similarly but with 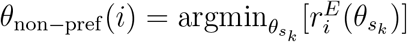.

In the experimental data we considered learning-related response changes of putative pyramidal cells to the vertical and angled grating corridors (see Khan et al. for how cells were identified). For each cell, we computed the difference in its response to the vertical and angled stimuli both pre- and post-learning 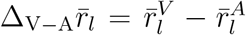 (where *l* = pre, post). We also computed the change in response to the vertical and angled stimulus over learning 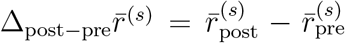 (where *s* = *A, V*). We then took the mean and variance of 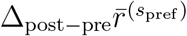 over all pyramidal cells which passed a set of inclusion criteria (where 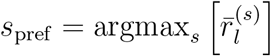 is the preferred stimulus of the cell). The inclusion criteria were as follows: the cell had a significant preference for one of the vertical and angled stimuli both before and after learning (defined as *p* < 0.05 under a Wilcoxon rank-sum test on the responses on vertical vs angled trials); the preferred stimulus *s*_pref_ was the same before and after learning. These criteria were necessary to avoid confounds relating to regression to the mean. The same analysis was performed for the non-preferred stimulus, in this case using 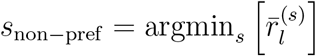.

We computed the average response SNR of individual E and I cells in both the model and data (Supplementary Figure 6E, F). The method for computing E and I response SNR in the network models is described in the above section. Quantification of mean SNR of individual pyramidal and parvalbumin cells was similar, and has been reported in Khan et al. (2018).

## Supplementary Figures

**Supplementary Figure 1.**
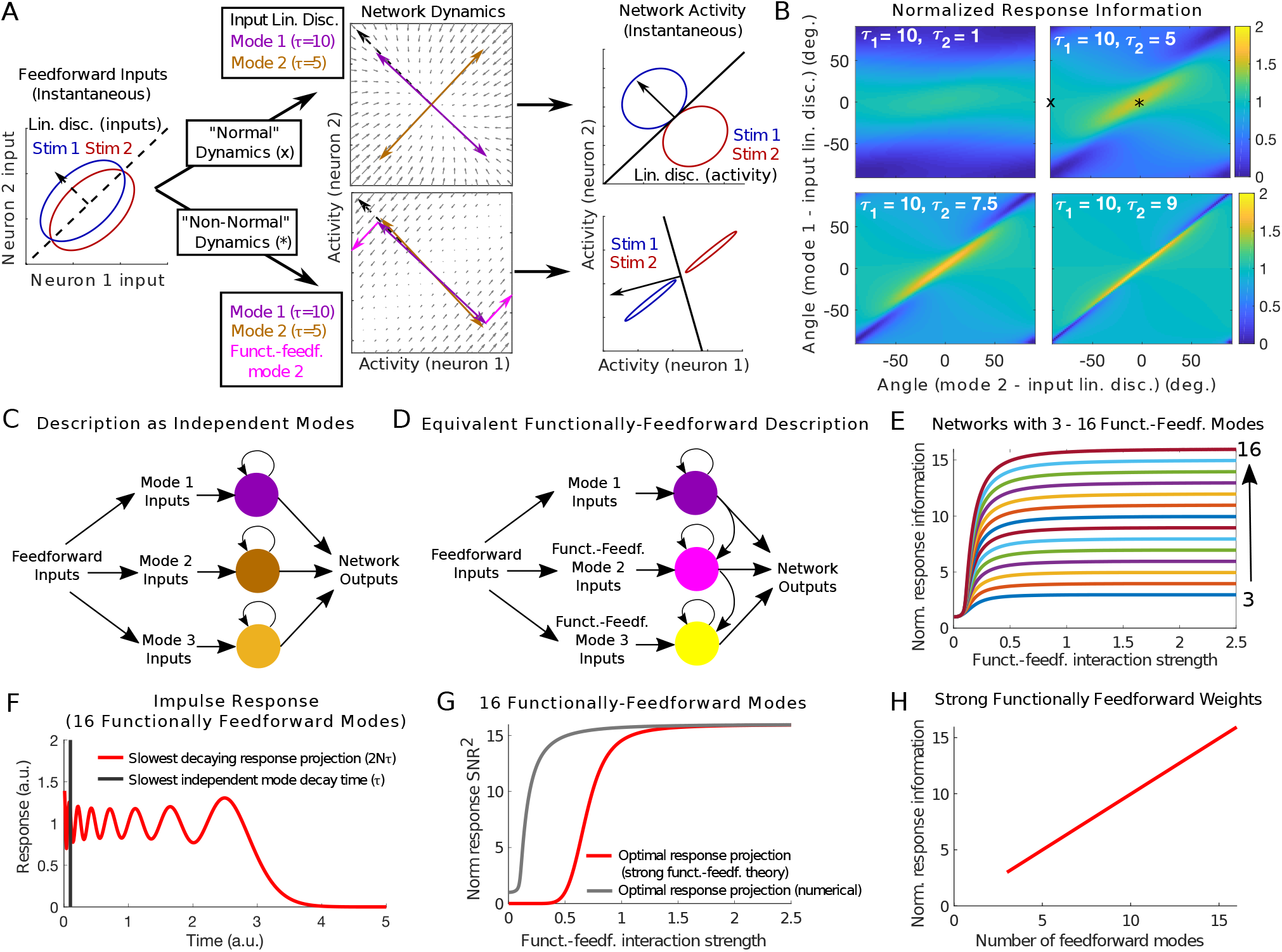
Non-normal dynamics can increase response information through functionally-feedforward temporal integration of the optimal input discriminant. A: Integration of feedforward input through normal and non-normal dynamics. Left: Distributions of instantaneous feedforward input for two stimuli and their linear discriminant. Middle: Recurrent dynamics around an input-driven fixed point. Non-normal dynamics can be described by either independent modes or functionally-feedforward modes (Schur decomposition or Jordan normal form; see panels C, D). Right: Distributions of instantaneous network activity following integration of feedforward input. B: Response information depends on the time constants and the activation patterns of modes. x and * are the parameters for the two example networks shown in A. Response information is normalized by the maximum information achievable in a normal network with the same time constants. Maximum response information occurs when both modes are aligned to the input discriminant and have similar time constants. C, D: Characterization of network dynamics by independent modes (eigen-vectors) or “functionally-feedforward” modes (e.g., Schur decomposition). Both are valid descriptions of the dynamics, but functionally-feedforward modes reveal non-normal integration more clearly. E: Response information for networks with varying numbers of functionally-feedforward modes and strength of functionally-feedforward interactions. Information is maximized in networks with strong functionally-feedforward dynamics and grows with the number of modes. F: Response of a strong functionally-feedforward network to a pulse of input. Black line shows the decay time constant of individual modes and red trace shows the time course of the most slowly decaying projection of network output. G: Squared SNR of two projections of network outputs. Red shows the optimal projection derived analytically assuming infinitely strong functionally-feedforward weights. Gray curve shows the optimal projection computed numerically for finite weights. H: Response information increases linearly with number of functionally-feedforward modes.

**Supplementary Figure 2.**
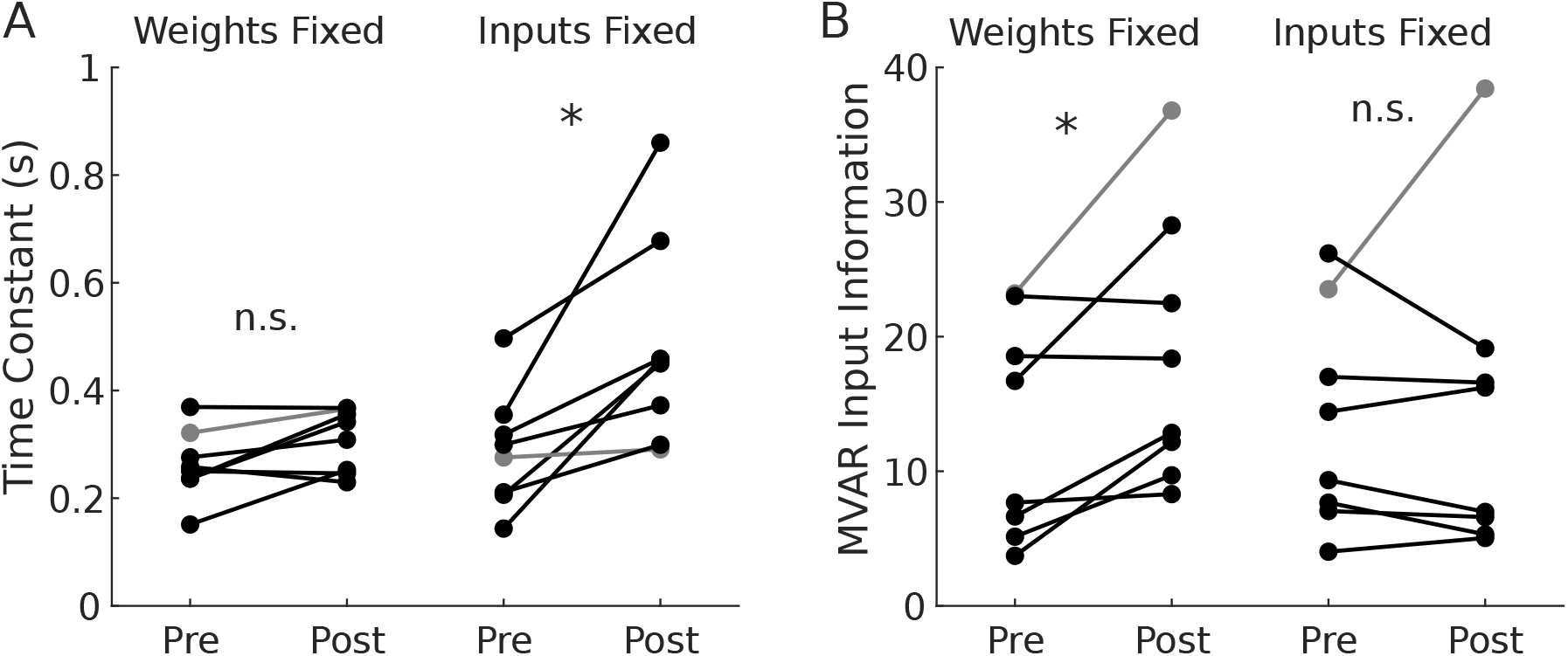
Improvements in temporal integration rely of reorganization of interaction weights but not stimulus-related input. A: Time constant of response to input along linear discriminant for an MVAR model in which interaction weights or stimulus-related input was constrained to be the same before and after learning. Gray line shows mouse whose time constant decreased over learning when all parameters were free (see Figure 3E, F, I). B: Information in stimulus-related input to MVAR model. Input information increased when weights were fixed, but not when input was fixed (note that input information could in principle improve through altered residuals even when mean input is held fixed).

**Supplementary Figure 3.**
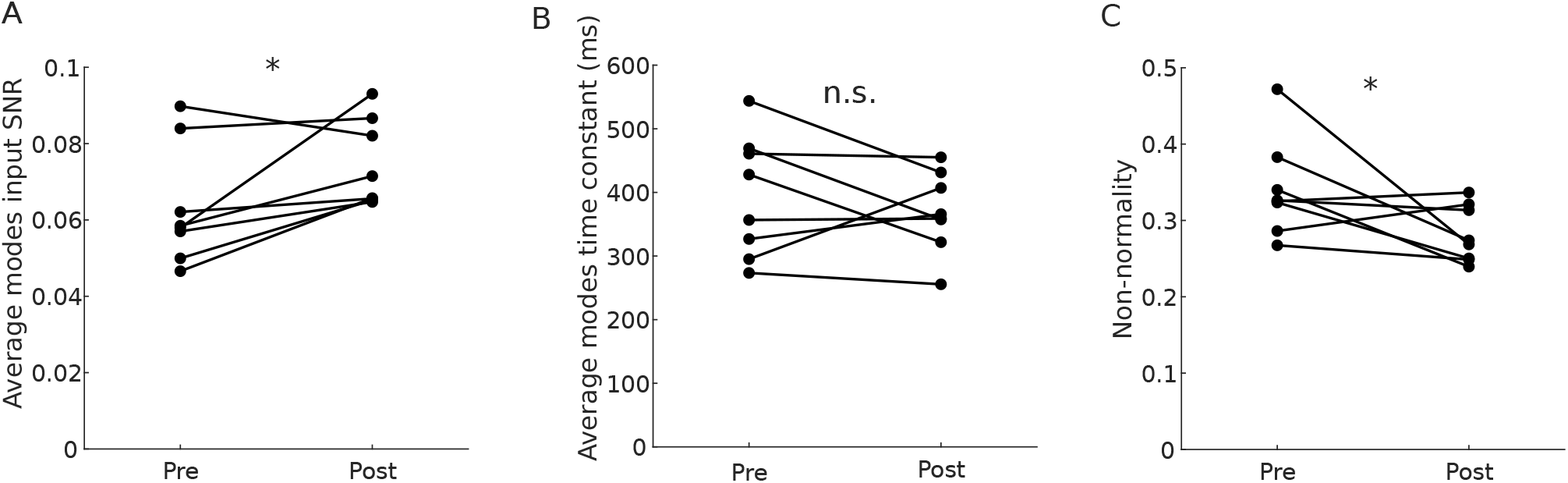
Individual mice show an increase in alignment of modes with the input linear discriminant, no increase in decay time constants, and a decrease in non-normality. A: Average over modes’ normalized input SNR, shown for each mouse pre- and post-learning. B: Average over modes’ time constant for each mouse. C: Non-normality of interaction weight matrices for each mouse pre- and post-learning.

**Supplementary Figure 4.**
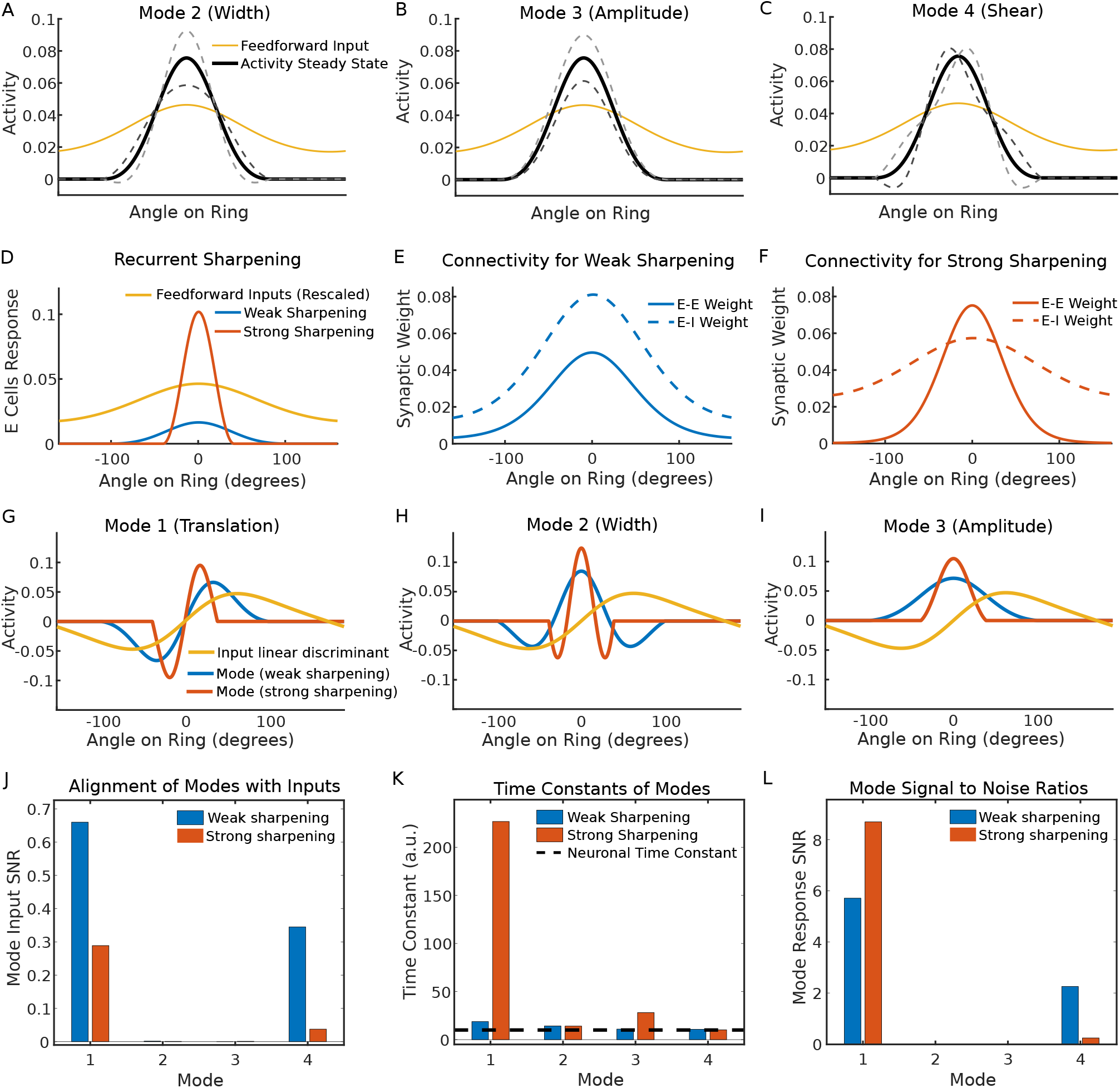
Uniform recurrent sharpening of sensory input reduces alignment of the slowest dynamical mode with the input linear discriminant. To test whether recurrent sharpening can explain the findings of the MVAR model, we examined the changes in the four slowest modes as connectivity was varied. A-C: Response steady state and perturbation along the 2nd-4th most slowly decaying modes in the E-I ring model (as in Figure 6B). D: Response of two networks to the same feedforward input, yielding weak and strong sharpening respectively. E, F: Patterns of network connectivity that induced the weak and strong sharpening of responses shown in D. Narrower E-E weights and/or broader E-I weights caused sharpening to increase (see Supplementary Figure 5 for a more comprehensive illustration). G-I: The activation patterns **m** of the three most slowly decaying modes, each overlaid with the input linear discriminant. In both networks, the translation mode was best aligned to the input discriminant and decayed most slowly. However, increased sharpening reduced the translation mode with the input discriminant (panel G, less overlap between the red and yellow curve than between cyan and yellow). J-L: SNR of feedforward input projected onto each mode (J), the time constant for each mode (K) and the SNR of network output along each mode (L). Although the decay time constant of the translation mode increased (panel K) and generated an increase in response SNR (panel L), these improvements are nonetheless inconsistent with the unchanged time constants and increased input SNR observed over learning in the MVAR model (Figure 5A, C).

**Supplementary Figure 5.**
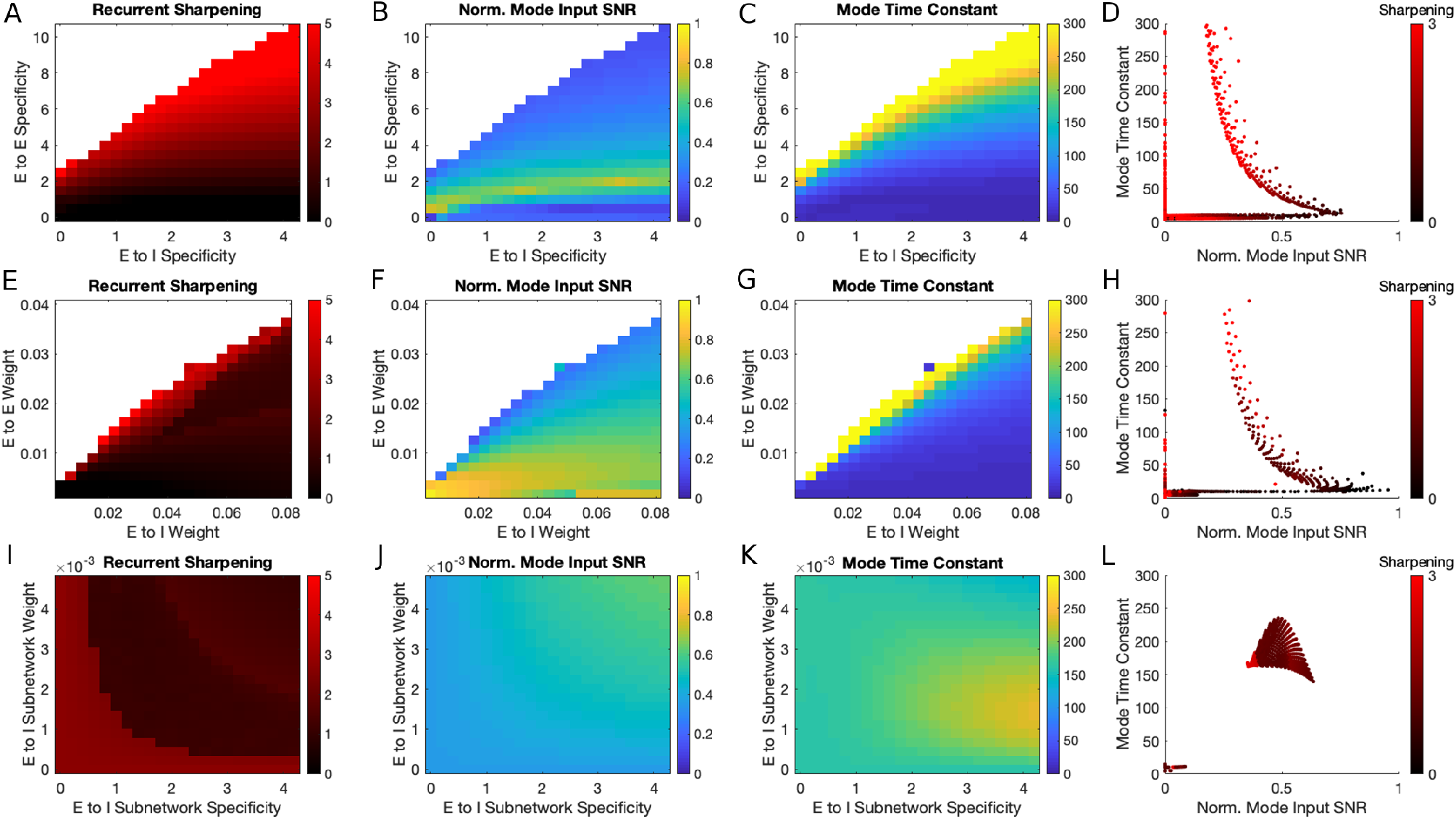
Parameter sweeps of excitatory-excitatory and excitatory-inhibitory synaptic weights. A: Degree of recurrent sharpening in networks with varying specificity (concentration around ring) of E to I and E to E weights. White denotes unstable networks (global instability or oscillation about an unstable fixed point). B: Normalized SNR of feedforward input projected along best mode (mode with greatest input SNR). C: Time constant of the mode shown in B. D: Modes pooled across networks shown in A-C (all modes pooled across all networks). For these uniform connectivity changes, time constants and normalized input SNRs covaried across networks and were largely constrained to lie on a 1-dimensional curve. For modes with decay time constants significantly greater than single-neuron time constants (here, 10), increases in normalized input SNR were consistently accompanied by decreases in time constant, in contrast to the stability of time constants with increased input SNR observed in the MVAR model. Although small increases in normalized input SNR with fixed time constant were possible (as evidenced by horizontal scatter about the main curve), these relied exclusively on a simultaneous reduction in the specificity of E-E and E-I synaptic weights and required fine-tuning of parameters to achieve. E-H: As in A-D but varying the magnitude of E to E and E to I weights across networks. I-L: As is A-D, but for networks with an E to I subnetwork of varying specificity and magnitude. These non-uniform connectivity changes yielded a fundamentally different relationship between mode time constant and input SNR, such that input SNR could be increased without altering decay time constants parameters by increasing the strength and tuning of the E-I subnetwork, with a wide range of connectivity parameters achieving the desired result.

**Supplementary Figure 6.**
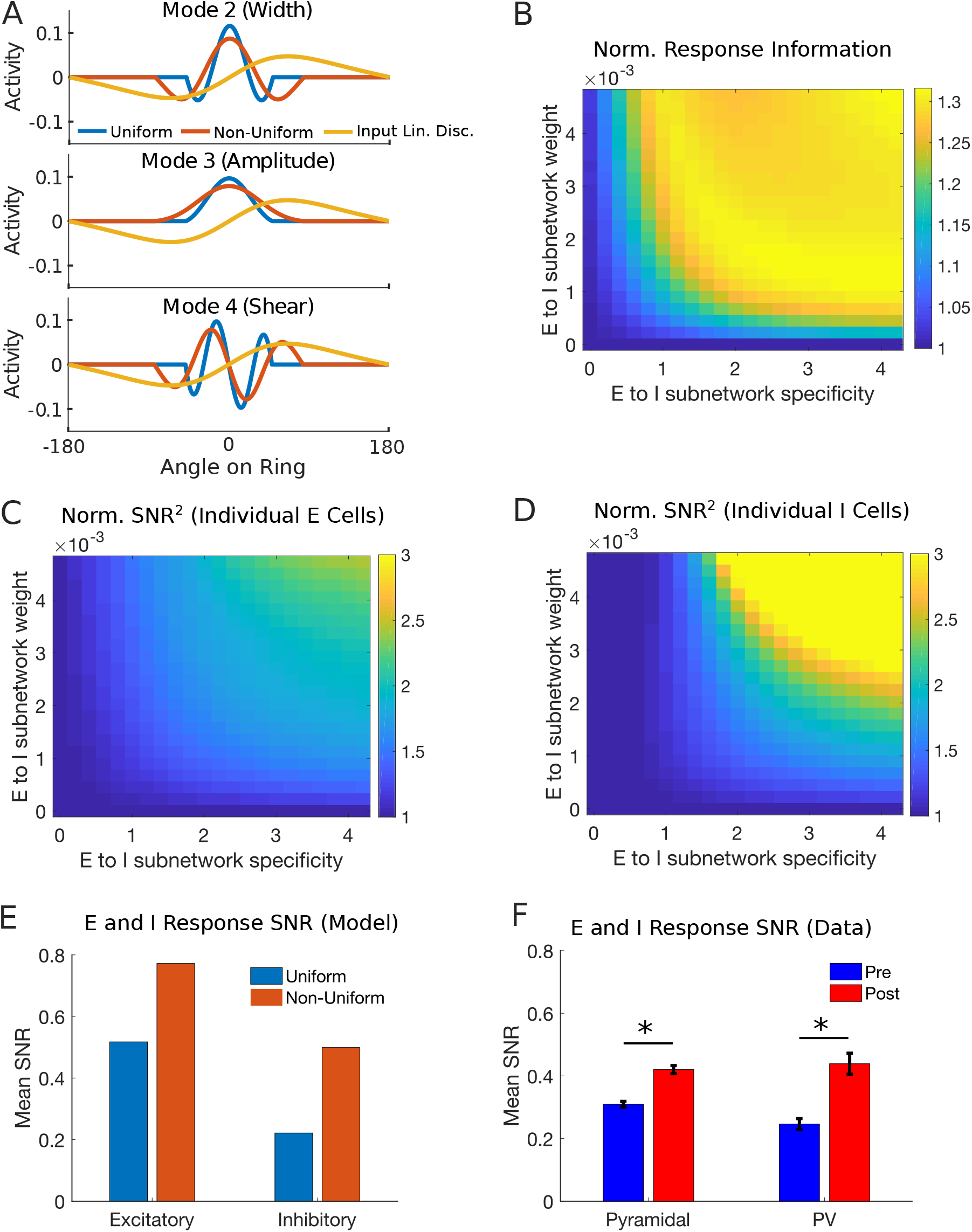
Modes and response information for networks with non-uniform connectivity. A: Activation patterns **m** for modes 2-4 in the uniform and non-uniform networks shown in Figure 6D. B: Linear discriminability of the two stimuli shown in Figure 6I, for networks with varying subnetwork strength and specificity (information normalized by value for uniform network). C, D: Average squared SNR of excitatory and inhibitory responses (normalized by value for uniform network). F, G: Average SNR of excitatory and inhibitory responses for the uniform and non-uniform network (unnormalized). G: Average SNR of excitatory (pyramidal) and inhibitory (PV) responses for the pre- and post-learning data.

## Supplementary Mathematical Note

### Notation

We use bold-face lower case letters for column vectors and non-bold upper case letters for matrices. Superscript *T* denotes a (vector or matrix) transpose; *x*_*i*_ or (**x**)*i* denotes the *i*th element of vector 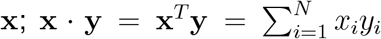 denotes an inner (dot) product of vectors; **xy**^*T*^ denotes an outer product of vectors with (*ij*)th element = *x*_*i*_*y*_*j*_; 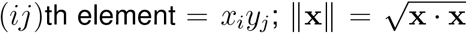 denotes the Euclidean vector norm; 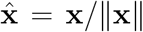 denotes a unit vector; 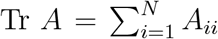 denotes the trace of an *N* × *N* matrix *A*; *I* denotes the identity matrix; we make use of the shorthand notation for the transpose of a matrix inverse *X*^−*T*^ =(*X*^*T*^)^−1^ = (*X*^−1^)^*T*^ ; ⟨**x**⟩ denotes the ensemble average of **x** (or time-average for ergodic variables); *δ*_*ij*_ denotes the Kronecker delta symbol and *δ*(*t*) denotes the Dirac delta function.

### Signal Processing Analysis

In this section we derive the results of Figure 1 in the main text. We consider a simplified model describing the sensory input to a network of neurons upon presentation a stimulus. Under the assumptions of this simple model, we derive the optimal method to discriminate a pair of stimuli based on observations of the network input. We also derive the performance of a more general class of suboptimal discrimination functions which we will later show are relevant to the way in which recurrent network dynamics act on the sensory input. This signal processing analysis places an upper bound on the possible discrimination performance of any network receiving such sensory input, specifies the mathematical operations a network must apply to its input in order to achieve this upper bound, and shows how suboptimal integration can be understood in terms of information loss both instantaneously and over time. In the sections that follow we use the results of this analysis to interpret the behavior of recurrent networks integrating such sensory input.

We consider a network of *N* neurons receiving sensory input **u** ∈ ℝ^*N*^ generated from a stimulus *s*. In the scenario we consider, one of two stimuli *s* ∈ {*s*_1_, *s*_2_} may be presented, each of which generates a time-series of sensory input **u**(*s, t*) drawn from a different distribution *p*(**u**|*s*). We assume that network input on any given trial consists of a time series **u**(*s, t*) = **g**(*s*)+*η*(*t*) with time-independent but stimulus-dependent mean **g**(*s*) and additive, stimulus-independent, multivariate normal noise *η*(*t*) ∼ *N*(**0**, Σ_*η*_) with ⟨*η*(*t*) ⟩ = 0 and ⟨*η*(*t*)*η*^*T*^ (*t*) ⟩ = Σ_*η*_. We wish to infer the identity of the stimulus *s* having observed a single realization of such a time series **u**. This can be achieved optimally by maximizing the posterior probability *p*(*s*|**u**) over the two stimuli.

We first consider how the two stimuli can be discriminated given an observation of network input **u**_0_ at a single time sample *t*_0_. In this case, the most probable stimulus *s* given the input vector **u**_0_ can be found using linear discriminant analysis (LDA), i.e. by taking a linear projection of the input vector **w** · **u**_0_ and comparing this to a threshold *c*. To see this, note that 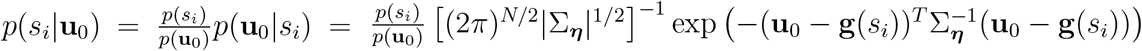, which gives 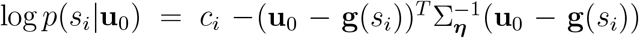 where *c*_*i*_ is a constant with respect to **u**_0_. Thus, 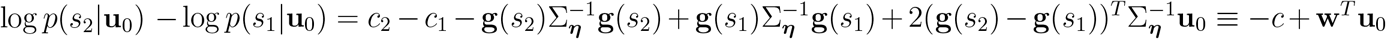, where we have absorbed all constant terms into a single scalar *c* and defined the projection vector 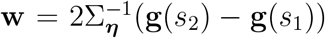. Therefore, the most probable stimulus given the observed input vector **u**_0_ is found by asking whether **w**^*T*^ **u**_0_ ≶ *c* (i.e., if **w**^*T*^ **u**_0_ > *c* then *s* = *s*_2_ is more probable, whereas if **w**^*T*^ **u**_0_ < *c* then *s* = *s*_1_ is more probable). The projection vector **w** is known as the linear discriminant, and can be understood as the vector which is normal to the hyperplane separating the two stimulus input distributions. The constant *c* determines the location of that hyperplane. Note that **w** and *c* can be rescaled by an arbitrary scalar constant without altering the decision rule.

We next consider how stimuli can best be discriminated when network input is observed sequentially in time. When statistically independent inputs **u**(*t*) are observed at a set of times *t* ∈ 𝒯 (a continuous interval or discrete samples), the optimal solution is to perform a time-averaged LDA using the decision rule **w** · ⟨**u**(*t*)⟩_*t*∈*t*_ ≶ *c*. Here, ⟨·⟩_*t*∈*t*_ is the sample mean over the set of time samples and **w**, *c* are the same quantities as in the single time sample case. This result follows directly from the single time sample case and the fact that 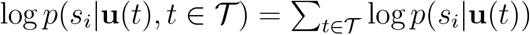 for statistically independent samples.

An intuitive way to understand this time-averaged LDA solution is to search for the linear projection **n** ∈ ℝ^*N*^ and temporal filter *f* (*t*) which, when applied jointly to the input time series **u**(*s, t*), generate the scalar output with the greatest signal to noise ratio with respect to the two stimuli to be discriminated. In the case of a continuous time series of length *T*, i.e. *t* ∈ [0, *T*], we denote the scalar output of such an operation as 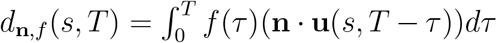. The signal to noise ratio of *d*_**n**,*f*_ (*s, T*) is defined as:

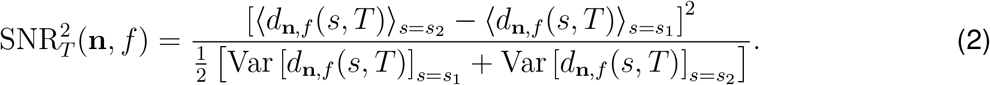

Provided that **u**(*s, t*) has Gaussian statistics, *d*_**n**,*f*_ (*s, T*) is a normally distributed random variable under each stimulus *s*. Moreover, assuming stimulus-independent input covariance, the variance of *d*_**n**,*f*_ (*s, T*) is independent of *s*. As a consequence, the above signal to noise ratio is sufficient to determine stimulus discrimination performance of an optimal observer receiving the scalar output *d*_**n**,*f*_ (*s, T*) (in particular, *p*(correct) = Φ(SNR_*T*_ /2) where Φ is the cumulative function of the standard normal distribution). The solution derived above by maximizing the posterior probability over *s* corresponds to setting 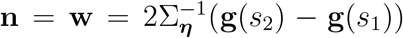. We rederive this optimal solution below through maximization of the above SNR. As we will show, using a different projection vector **n** or temporal filter *f* reduces the signal to noise ratio (except for scaling of *f* or **n**, which has no effect). Thus, the linear discriminant vector **w** can also be understood as the vector which maximizes the signal to noise ratio of the projected input.

We now derive the optimal choice of **n**, *f* and quantify the performance of both optimal and sub-optimal choices under the assumption of temporally uncorrelated Gaussian input noise. In this case, the influence of **n** and *f* on the signal to noise ratio of the scalar output *d*_**n**,*f*_ (*s, T*) takes on a particularly simple form. In particular, we then have ⟨*η*(*t*)*η*^*T*^ (*t*′)⟩= Σ_*η*_*δ*(*t* − *t*′), so that 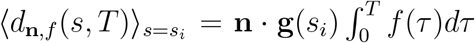 and 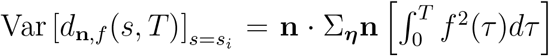. Defining Δ**g** = **g**(*s*_2_) − **g**(*s*_1_), the output signal to noise ratio is then given by:

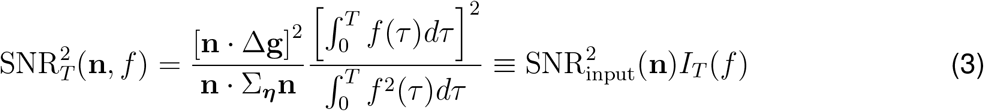

where 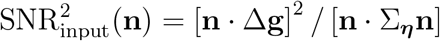 is the signal to noise ratio of the instantaneous input projected along **n** and 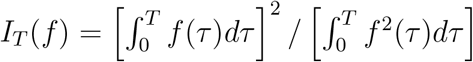 is a temporal integration factor. Thus, the total signal to noise ratio factors into an instantaneous term and a temporal term. We can therefore proceed to maximize each of these two factors in turn with respect to **n** and *f* respectively. To do so, we apply the Cauchy-Schwarz inequality to derive two inequalities, 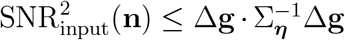 and *I*(*f*) ≤ *T*. To see how the first inequality arises, note that 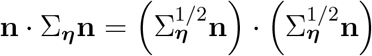, while by Cauchy-Schwarz 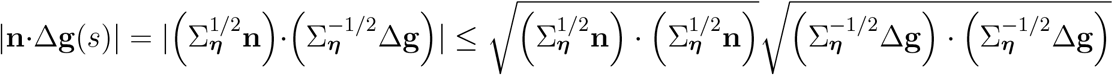. Inserting these into the definition of 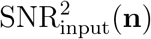 and cancelling terms in the numerator and denominator gives the desired inequality. The second inequality follows in a similar fashion: the integral Cauchy-Schwarz inequality gives 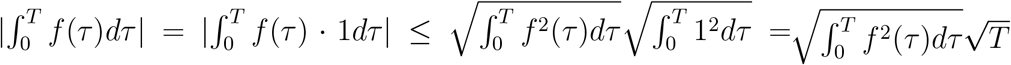 which can be inserted into the definition of *I*_*T*_ (*f*) to arrive at the desired result. It can easily be verified that these upper bounds are achieved when *f* (*t*) = *α* and 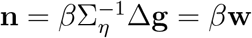 for any pair of constants *α, β*. Thus, we have arrived at the same optimal solution for stimulus discrimination using two different methods: first, by maximizing the posterior probability of the stimulus given the observed network input; second, by maximizing the signal to noise ratio obtained by linear projection and temporal filtering of the network input.

Several conclusions can be drawn from this analysis. First, for invertible Σ_*η*_, the information available to a decoder of network input over a time window *T* is finite and the sources of information loss can be factored into an instantaneous term SNR_input_ and a temporal term *I*_*T*_ (*f*) (note that further sources of information loss may occur when different functions than those considered here are applied to the network input, as we will see when we study recurrent networks). Moreover, even in the limit of infinite time, the information available to decoder with finite timescales of temporal integration remains finite due to the loss of previously integrated information over time (i.e., if lim_*T*→∞_ *I*_*T*_ (*f*) < ∞). As we have shown, the optimal solution for discriminating pairs of stimuli given an observed time series of network input is to project that network input onto the direction carrying the most information instantaneously, and then to integrate that projection using a sufficiently long time constant in order to avoid loss of previously integrated information (i.e., using a choice of *f* such that *I*_*T*_ (*f*)/*T* ≈ 1). In the following analysis of information transmission through recurrent networks, we will focus on the information contained in the output of networks with finite dynamical time constants following integration of sensory input over a long period of time.

### Analysis of Fisher Information in Recurrent Networks

We next quantify the capacity of an optimal observer to discriminate stimuli based on observations of the output of a recurrent network which receives the sensory input described in the previous section. We analyze the transformation of noisy sensory input by a recurrent network of *N* nonlinear units governed by the following dynamics:

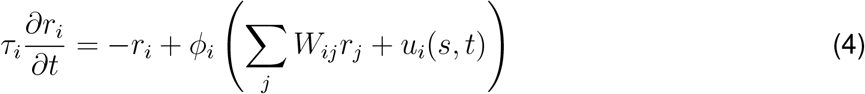

where *r*_*i*_ represents the firing rate of neuron *i, τ*_*i*_ is its time constant, *ϕ*_*i*_ is its input-output nonlinearity (or transfer function), *W*_*ij*_ is the synaptic weight from neuron *j* to neuron *i* and *u*_*i*_(*s, t*) = *g*_*i*_(*s*)+*η*_*i*_(*t*) is the feedforward input to neuron *i* at time *t* given a sensory stimulus *s*. As before, inputs are defined as having additive, multivariate Gaussian, temporally uncorrelated, stimulus-independent noise *η*(*t*).

Rather than deriving the signal to noise ratio for two discrete stimuli as above, we will derive the Fisher Information of network responses **r** with respect to a continuous one-dimensional stimulus *s*. The Fisher Information places a lower bound on the variance of any unbiased estimator of *s* from **r**. For responses following a multivariate normal distribution, the Fisher Information is given by 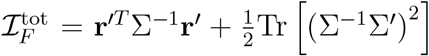, where 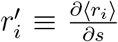 is the slope of the tuning curves with respect to *s*, Σ = ⟨(**r** − ⟨**r**⟩) (**r** − ⟨**r**⟩)^*T*^ ⟩ is the covariance of network responses under that stimulus and 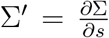 is the change in response covariance as the stimulus is changed. When Σ is stimulus-dependent, achieving the precision of stimulus discrimination set by the Fisher Information requires a quadratic decoder of neural activity (Shamir and Sompolinsky, 2004; Yang et al., 2020). We focus instead on the linear Fisher Information ℐ_*F*_ = **r**^′*T*^ Σ^−1^**r**′ following previous studies (Seriès et al., 2004; Beck et al., 2011; Moreno-Bote et al., 2014). In addition to being analytically tractable, the linear Fisher Information has several theoretical advantages. First, even for networks in which the optimal decoder is quadratic (or otherwise nonlinear), the linear Fisher Information describes the optimal local linear decoder of small changes in the stimulus based on network responses (Seriès et al., 2004; Beck et al., 2011; Kafashan et al., 2021). Second, the linear Fisher Information places a bound on the precision of an optimal linear estimator even for non-Gaussian response distributions, whereas the quadratic term holds only for Gaussian statistics (Yang et al., 2020; Kafashan et al., 2021). Third, the linear Fisher Information has a natural relationship to linear discriminant analysis, in particular ℐ_*F*_ Δ*s*^2^ ≈ Δ**r**^*T*^ Σ^−1^Δ**r** for sufficiently small Δ*s*, which allows us to relate our findings back to the two-stimulus discrimination task studied experimentally in the main text and above in our signal processing analysis. Fourth, the linear Fisher Information can be understood as a signal to noise ratio, much as in our above signal processing analysis. In particular, the linear Fisher Information is the SNR of **w**^*T*^ **r**, where **w** = Σ^−1^**r**′ is the linear discriminant vector for discriminating infinitesimal changes in *s* based on network output **r**.

In order to evaluate the linear Fisher Information of the output of a recurrent network, we next derive expressions for the tuning curve derivatives **r**′ and response covariance Σ for networks obeying the dynamics of Equation (4) and driven to stationary state.

#### Tuning Curve Slopes and Response Covariance

The linear Fisher Information of the output of a recurrent network **r** depends on two quantities: the tuning curves with respect to the stimulus 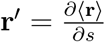, and the response covariance Σ = ⟨ (**r** − ⟨**r**⟩)(**r** − ⟨**r**⟩)^*T*^⟩. To derive expressions for these, we will rely on two approximations: first, we linearize the system about a stimulus-evoked fixed point; second, we compute the statistics of the stationary state response of the linearized system.

To estimate the tuning curve derivatives 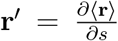, we differentiate the noise-free fixed points of the network with respect to the stimulus. To do so we set 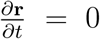 and *η* = 0 and then differentiate both sides of Equation (4) with respect to *s*. On performing this calculation, we obtain **r**′_*SS*_(*s*) = −*J*^−1^(*s*)Φ′(*s*)**g**′(*s*), where **r**_*SS*_(*s*) = *ϕ* (*W* **r**_*SS*_(*s*) + **g**(*s*)) is the noise-free steady state response, *J*(*s*) = Φ′(*s*)*W* − *T* ^−1^ is a matrix of effective interaction weights and 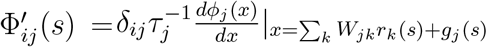 is a diagonal matrix quantifying the sensitivity of each neuron to small changes in its input (both feedforward and recurrent). Note that this result involves an approximation: we have replaced the average stationary state response of the stochastic system ⟨**r**⟩ with the fixed point of the noise-free system **r**_*SS*_. The accuracy of this approximation depends on the nonlinearity near the fixed point and on the magnitude of the noise. Note that while we did not explicitly linearize in order to obtain this solution, an identical result is obtained by by first linearizing the network dynamics about the noise-free fixed point, computing the mean response of the noise-injected linearized system at stationary state, and then differentiating this with respect to the stimulus. This is the approach we next take in order to obtain an approximation for the response covariance.

To derive the response covariance within the linearized stationary state approximation, we first linearize Equation (4) about the fixed point **r** = **r**_*SS*_(*s*) by applying a first order Taylor expansion for small fluctuations *δ***r** about the fixed point **r**_*SS*_, i.e. **r** = **r**_*SS*_ + *δ***r** with ‖*δ***r**‖ ≈ 0. This gives the following approximation to the dynamics:

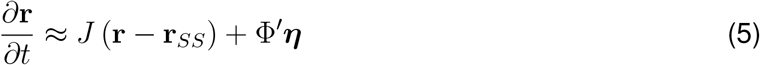

where *J* and Φ′ are as defined above. Equation (5) describes a multivariate Ornstein-Uhlenbeck process, and has the general solution:

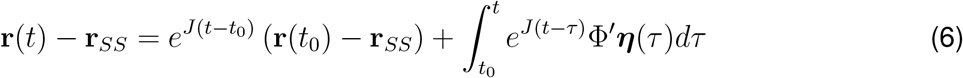

for any initial condition **r**(*t*_0_), where *e*^*X*^ is the matrix exponential function. Provided the fixed point is stable (i.e., all eigenvalues of *J* have negative real part) we can take the stationary state limit by letting *t*_0_ → −∞ to obtain:

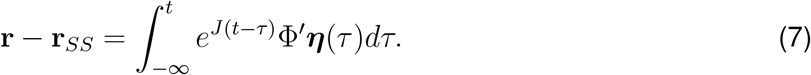

Assuming that input noise is temporally uncorrelated, i.e. ⟨*η*(*t*)*η*^*T*^ (*t*′)⟩ = Σ_*η*_*δ*(*t* − *t*′), the stationary-state response covariance Σ_*SS*_ = ⟨(**r** − **r**_*SS*_) (**r** − **r**_*SS*_)^*T*^ ⟩ is:

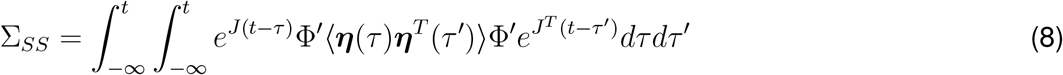

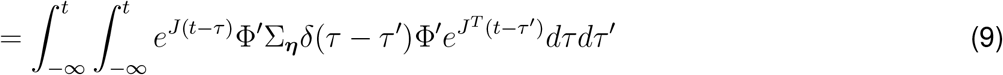

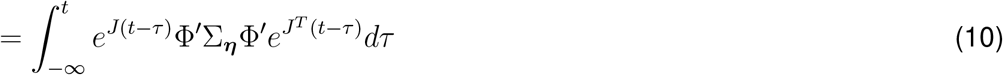

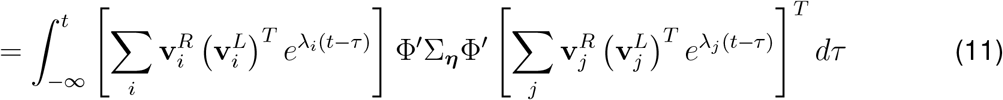

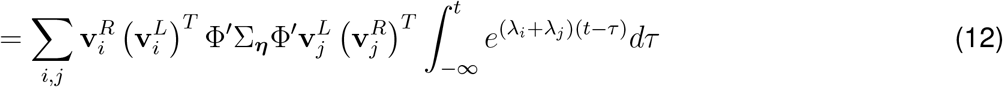

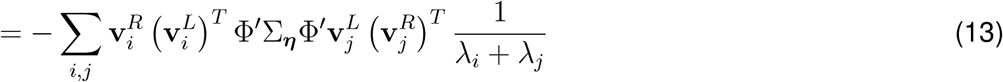

where we have made use of the eigendecomposition of the Jacobian 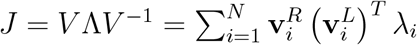 and of its matrix exponential 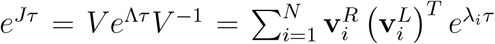. We use superscripts *L* and *R* to denote left and right eigenvectors, which are the rows of *V* ^−1^ and columns of *V* respectively. Note that the left and right eigenvectors do not in general form orthonormal bases, but do satisfy the orthogonality relations 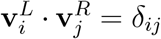. This orthogonality relation does not typically allow for both left and right eigenvectors to have unit length, because 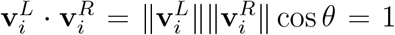. Where a choice of normalization is required, we choose to normalize left eigenvectors to unit length, in which case right eigenvectors typically do not have unit length. This convention for normalization is entirely arbitrary and is made for convenience only, reflecting the central role that left eigenvectors play in our theory. In the main text, we refer to the left eigenvectors as the mode activation patterns **m**, and we define their time constants as *τ* = −1/Re(*λ*). Note that the stationary state covariance also satisfies the Lyapunov equation *J*Σ_*SS*_ + Σ_*SS*_*J*^*T*^ + Φ′Σ_*η*_Φ′ = 0, which is well known in the control theory literature. This Lyapunov equation can be solved efficiently using numerical methods, but is less convenient when deriving the analytical results we present the following sections.

#### Relationship Between Eigen-Modes and Signal Processing Theory

With the results of the previous section in hand, we are now in a position to formulate a general expression for the Linear Fisher Information of the network response. Before doing so, however, we first show that the signal to noise ratio of the network output projected along any left eigenvector (i.e., mode) 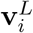 of the Jacobian *J* takes on a particularly simple form that is readily interpretable using the insights obtained from our earlier signal processing analysis. The linear Fisher Information can be understood as the signal to noise ratio of network output projected onto the linear discriminant vector for the network output, which in turn can be understood as the projection vector which maximizes this signal to noise ratio (as shown in our signal processing analysis). Thus, deriving an expression for the signal to noise ratio along any other projection (in this case, a left eigenvector) allows us to place a lower bound on the total information in the network response. The equations derived in this section form the basis for the results presented in Figure 2 of the main text, and motivate much of our analysis of the experimental data and network models presented in Figures 3-6.

To simplify the expressions which follow, we first make a change of variables 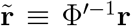 and 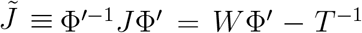. In this basis, Equation (5) becomes 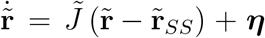, while 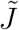 has eigenvalues 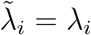 and eigenvectors 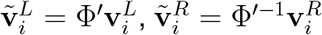. We can express the tuning curve derivatives as 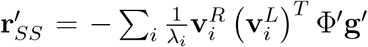. Then using the identity 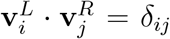, both Σ_*SS*_ and **r**′_*SS*_ can be expressed in the basis of left eigenvectors, which obtains:

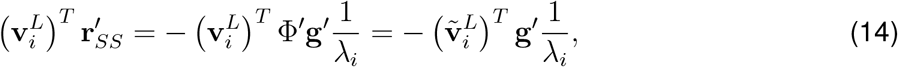

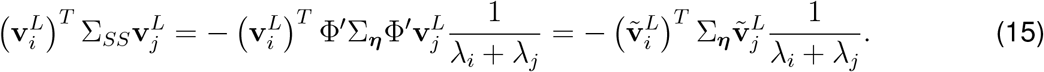

We can then calculate the signal to noise ratio of the instantaneous network response at stationary state, projected along any left eigenvector 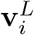:

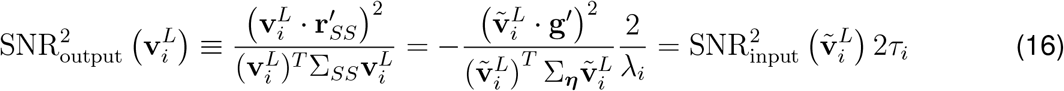

where we have defined *τ*_*i*_ = -1/*λ*_*i*_, under the assumption that *λ*_*i*_ ∈ ℝ (i.e., the mode is not oscillatory).

Equation (16) demonstrates that the SNR of network output following projection onto any left eigen-vector of *J* is equal to the SNR of network input projected along the corresponding left eigenvector of 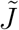, multiplied by the decay time constant of that eigen-mode and by a constant factor of 2. This result is identical to that obtained in our signal processing analysis, and can easily be derived from Equation (3) by setting 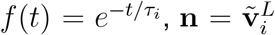, and taking *T* → ∞. The reason for this correspondence is that left eigenvectors implement exactly the linear projection and temporal filtering operations required for optimal stimulus discrimination, up to the minor caveat that the optimal (but biologically implausible) *f* (*t*) = 1/*T* is replaced with an exponential filter 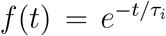. We can identify the scalar output *d*_**n**,*f*_ (*s, T*) from the signal processing analysis with the linear projection of the network response 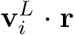. Equation (16) is the main result presented in Figure 2, where we considered a purely linear (rather than linearized) system, which slightly simplifies the result because 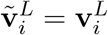.

It is important to emphasize that, while Equation (16) can be understood as a special case of our more general signal processing analysis (which allows for arbitrary filters *f* (*t*)), this result in fact relies on the unique properties of left eigenvectors. For example, a similar result is not obtained when projecting responses along right eigenvectors 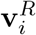. Indeed, there is a deeper reason that left eigenvectors exhibit this property. This result relies on two facts: first, network input along each left eigenvector is mapped onto network output along the corresponding right eigenvector; second, left eigenvectors are orthogonal to right eigenvectors 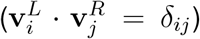. Together, these properties ensure that the network dynamics decouple into independent leaky integrators when projected onto left eigenvectors, in particular 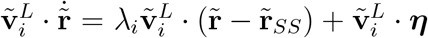 (and also 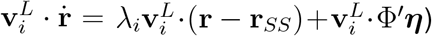. This decoupling into independent processes is a unique feature of the left eigenvector basis, and motivates the use of the word “modes” to describe them. This observation underscores an additional source of information loss in recurrent networks that was not apparent from our signal processing analysis - because recurrent networks map multiple different projections of their input onto any given projection of their output, they superimpose both relevant information and additional irrelevant noise within the same output projection, which reduces the signal to noise ratio. Left eigenvectors avoid this source of information loss by isolating a single projection of network input and preserving it along a single projection of the network output, allowing them to integrate input information optimally.

#### Linear Fisher Information at Stationary State

We now return to the problem of estimating the linear Fisher Information of the network response. The Linear Fisher Information is equal to the signal to noise ratio obtained after projecting network responses along their linear discriminant 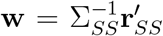. Because the linear discriminant is the projection which maximizes this signal to noise ratio, the linear Fisher Information will typically exceed the signal to noise ratio obtained following projection along any left eigenvector (Equation (16)). Inserting the expressions for tuning curve slopes and response covariance derived above into the equation for the linear Fisher Information, we obtain:

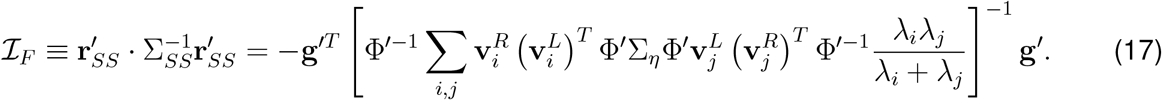

Using again the change of basis introduced in the previous section, this result simplifies to:

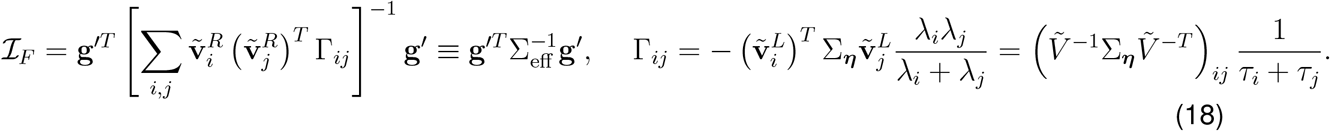

This equation provides intuition as to how the transformation of sensory input through the recurrent network shapes the information about the stimulus available in the network output. The linear Fisher Information of the instantaneous sensory input is 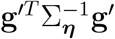, so that Σ_eff_ encapsulates the relationship between input and output information (the transformation of both input signal and noise by the network have been absorbed into this effective covariance). The coefficients Γ_*ij*_ have a natural interpretation as the effective covariance between network responses projected onto pairs of left eigenvectors, i.e. 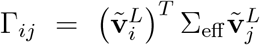 and 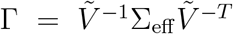. Moreover, these coefficients depend on the alignment of the corresponding pair of left eigenvectors with the sensory input covariance and also depend inversely on the timescale of integration along those eigenvectors *τ*_*i*_ + *τ*_*j*_ = − (*λ*_*i*_ + *λ*_*j*_) / (*λ*_*i*_*λ*_*j*_) (assuming the eigenvalues are real). Moreover, Γ is the solution to the Lyapunov equation 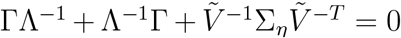, meaning it is the stationary state covariance of a system with injected covariance 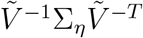 and dynamical evolution Λ^−1^. Similarly, the effective covariance follows the Lyapunov equation 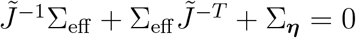.

The Fisher Information can be expressed compactly in matrix form as:

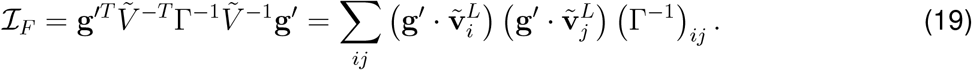

Unfortunately, this expression for Fisher Information is difficult to compute analytically except in certain special cases where Γ can be directly inverted, such as when Γ is a 2×2 matrix or a diagonal matrix. For a diagonal Γ we have:

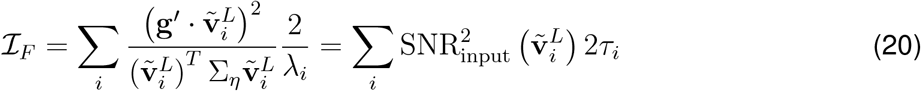

so that the Fisher Information in the network response is simply the sum of response SNRs along individual left eigenvectors. Although this case provides useful intuition, the assumption that Γ is diagonal places strong restrictions on the dynamics which may not be applicable to neural circuits, for example that the eigenvectors are orthogonal. For such networks (also known as “normal” networks), it can be seen that the solution which maximizes the linear Fisher Information in Equation (20) is to align the left eigenvector with the longest decay time constant *τ*_*k*_ with the linear discriminant of the instantaneous sensory input, so that 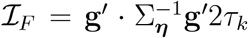 much as in our analysis of single eigen-modes.

#### Linear Fisher Information for Non-Normal Networks

Networks in which the eigenvectors of the Jacobian are not orthogonal are known as “non-normal” networks (Ganguli et al., 2008; Goldman, 2009; Murphy and Miller, 2009). We now study how non-normal network dynamics influence information integration and transmission. Our main finding is that non-normal dynamics can enhance the linear Fisher Information of network responses by a factor of up to *N* (the number of neurons in the network). These findings form the basis of the results presented in Supplementary Figure 1 of the main text. We note that closely related findings have been presented previously (Ganguli et al., 2008; Goldman, 2009). To arrive at these results, we first analyze the an arbitrary two-dimensional non-normal system, then use the optimal solution obtained in this 2-dimensional case to motivate a specific class of N-dimensional networks which achieve the desired N-fold improvement in information transmission.

To gain intuition into how non-normality of network dynamics affects linear Fisher Information, we perturb the solution obtained for the normal network adding a single pair off-diagonal elements Γ_*ab*_ = Γ_*ba*_ to Γ. This perturbed system corresponds a network in which only a two-dimensional plane exhibits non-normal dynamics, with the remaining eigenvectors forming an orthogonal basis. This system has effective covariance matrix 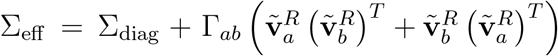, where Σ_diag_ is the effective covariance matrix for the unperturbed system. This covariance matrix can be inverted exactly using the Sherman-Morrison matrix inversion identity:

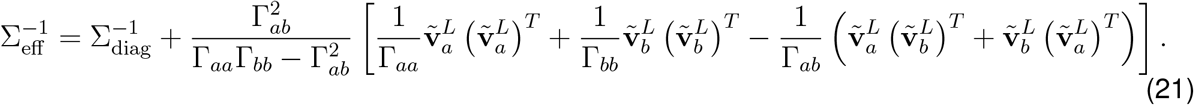

This result can then be used to obtain the linear Fisher Information of the perturbed system via Equations (18, 20):

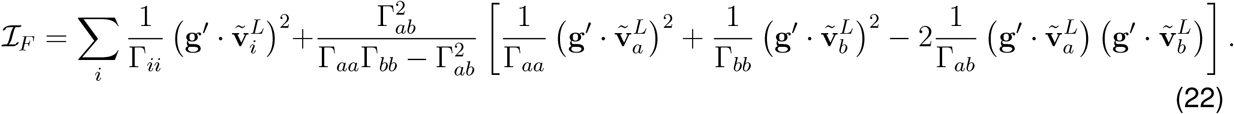

By rearranging this expression, we can make explicit the information contained in the non-normal plane of dynamics (given in the second term below):

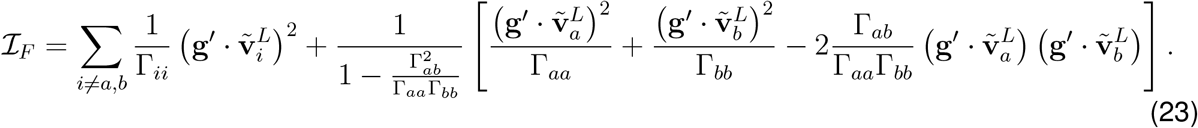

To understand how the non-normal component of the Fisher Information depends on the relative alignment of eigenvectors and their time constants we define 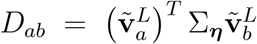, so that Γ_*ab*_ = *D*_*ab*_/(*τ*_*a*_ + *τ*_*b*_). We then introduce the two dimensionless quantities *β* = *τ*_*b*_/*τ*_*a*_ and 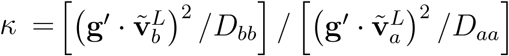. The term *D*_*ab*_ quantifies the degree of non-orthogonality of the eigenvector pair *a, b* (more precisely, the covariance of sensory input projected onto the pair of eigenvectors). *β* quantifies the relative time constants of the two eigen-modes, and *κ* quantifies the relative signal to noise ratio of sensory input projected onto the two left eigenvectors. Without loss of generality, we may assume that *τ*_*a*_ ≥ *τ*_*b*_, so that *β* ≤ 1.

Inserting these definitions into Equation (23) gives:

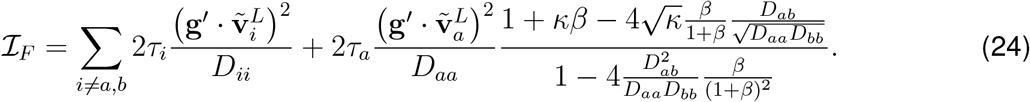

As *D*_*ab*_ → 0, the solution for the normal system is recovered (Equation (20)). However, if both *κ* → 1 and 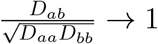 then the Fisher Information becomes 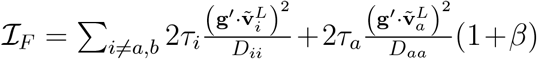. Then as *β* → 1 the Fisher Information becomes 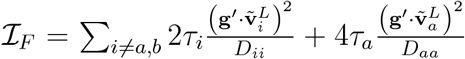. Taking this set of limits corresponds to the case where 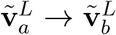 and *τ*_*a*_ → *τ*_*b*_. The linear Fisher Information is then maximized by setting 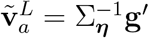, in which case both left eigenvectors in the non-normal plane are aligned to the input linear discriminant while all other left eigenvectors are orthogonal. The total response information for such a network is 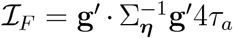, which is twice that achievable by any normal network whose longest time constant is *τ*_*a*_ (see Supplementary Figure 1B for a numerical validation of this result). It is noteworthy that the limit taken here yields a defective matrix 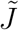, i.e. one which has fewer distinct eigenvectors than it has dimensions *N*. We next show that, by constructing a maximally-defective matrix, i.e. one which has just one eigenvector repeated *N* times, it is possible to achieve an *N*-fold improvement in linear Fisher Information relative to an optimal normal network.

To extend this two-dimensional example to the N-dimensional case, we construct a network in which non-normal dynamics produce an N-fold increase in response information. Motivated by our signal processing analysis, we search for cases in which there exists a pair of projections **w** of the neural response 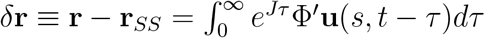 and **n** of the sensory input **u**(*s, t*) such that:

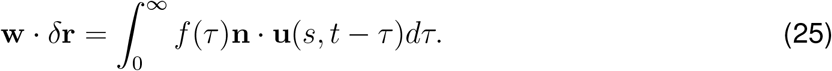

for some yet-to-be-determined function *f*(*t*). In such a case the SNR of network responses projected onto **w** is given by Equation (3) with *T* → ∞.

We can immediately identify one solution to Equation (25), which is 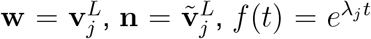. This recovers our single-eigenvector analysis. To construct a second case, we consider a network with *J*_*ij*_ = *λδ*_*ij*_ +*ωδ*_*i,j*-1_, which corresponds to a delay line in which units have decay time constants *τ*_*i*_ = −1/*λ* and feedforward weights *ω* (by feedforward, we mean that the weights are ordered along the delay line). It can be verified that this matrix has only one distinct eigenvalue *λ* and one distinct eigenvector (**v**^*L*^)_*i*_ = *δ*_*iN*_. Then 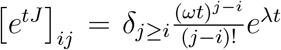 (as can be shown using the power series definition of a matrix exponential). Thus, Equation (25) becomes:

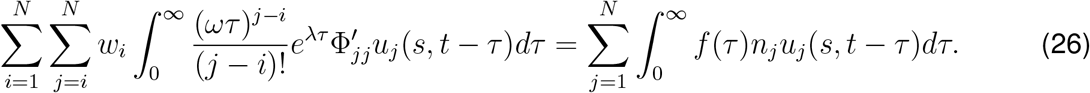

There does not in general exist an **n** and *f* which satisfy this equation, but in the limit *ω* → ∞ a solution exists because 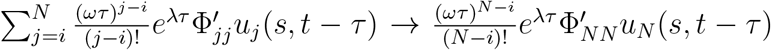. This gives the equation:

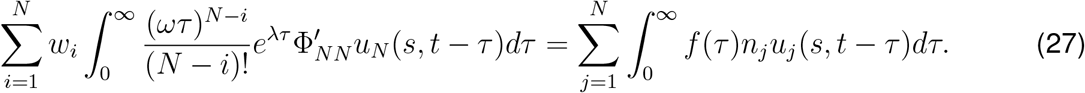

We can then identify a second solution to Equation (25), which is *n*_*i*_ = *δ*_*iN*_ and 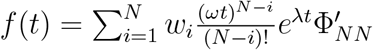. Thus, while we are free to choose any set of readout weights **w**, only the input to the *N*th neuron can be recovered from the output of such a network regardless of the readout weights we choose. In this case, the readout weights **w** determine the temporal filter *f* (*t*) applied to the *N*th neuron’s input, with different choices of **w** allowing different functions of the input history to be recovered.

Having identified this solution, we next proceed to maximize the SNR of responses along **w**. To optimize response SNR along **w**, we need to maximize both SNR_input_(**n**) and *I*_∞_(*f*) as defined in Equation (3). *I*_∞_(*f*) can be maximized by choosing the appropriate readout weights **w** as follows:

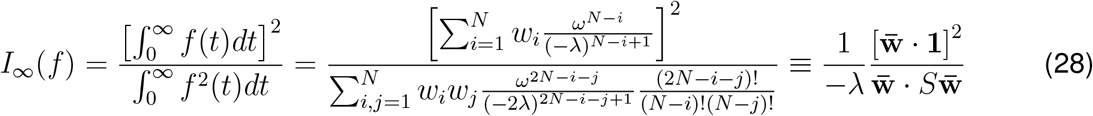

where we have defined 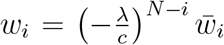 and 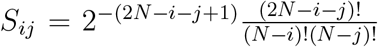 and **1** is a vector of ones. The Cauchy-Schwarz inequality then yields 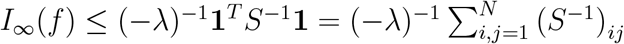, with the upper bound achieved when 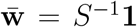. We find numerically that 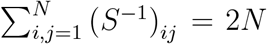, so that *I*_∞_(*f*) = (−*λ*)^−1^2*N*, revealing an N-fold increase in temporal integration through non-normal dynamics (because *λ* is the only eigenvalue of *J*, a normal network could obtain at best *I*_∞_(*f*) = 2(− *λ*)^−1^). Supplementary Figure 1F shows the temporal filter *f* (*t*) that results from this choice of weights when *N* = 16.

We now ask how to maximize the second factor in our signal processing analysis, SNR_input_(**n**). Because the input projection integrated by the above network is *n*_*i*_ = *δ*_*iN*_, SNR_input_(**n**) is maximized when the linear discriminant of sensory input is aligned to the *N*th element of the delay line. However, orthogonal transformations of this delay line, *J* → *U JU* ^*T*^ with *U* ^*T*^ = *U*^−1^, change the projection of sensory input integrated by the network as **n** → *U* **n**, but do not otherwise affect the results. Thus, SNR_input_(**n**) is maximized by rotating the delay line in neural space so that **n** aligns with the linear discriminant of sensory input, while *I*_∞_(*f*) is maximized by the appropriate choice of read-out weights **w** as described in the preceding paragraph (which must also be rotated, **w** → *U* **w**). This rotated delay line corresponds to a “functionally feedforward” dynamic (Goldman, 2009) and the integrative properties of such delay line architectures have been studied previously (Ganguli et al., 2008). The Jacobian *J* introduced here is a defective matrix, i.e. it has only one eigenvector 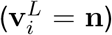 and one eigenvalue (*λ*), and therefore is consistent with the result of the two-dimensional case in which information increases when eigenvectors become more aligned and eigenvalues simultaneously become more similar. Moreover, the optimization of SNR_input_(**n**) requires that this left eigenvector is aligned to the input linear discriminant, demonstrating that the optimal non-normal network is one in which all left eigevectors are aligned to the input linear discriminant and have identical time constants. Supplementary Figure 1E-H show the response information computed from networks with varying number of units *N* and feedforward weight *ω*.

